# Reversing the decline of threatened koala (*Phascolarctos cinereus*) populations in New South Wales: Using genomics to define meaningful conservation goals

**DOI:** 10.1101/2023.12.03.569474

**Authors:** Matthew J. Lott, Greta J. Frankham, Mark D.B. Eldridge, David E. Alquezar-Planas, Lily Donnelly, Kyall R. Zenger, Kellie A. Leigh, Shannon R. Kjeldsen, Matt A. Field, John Lemon, Daniel Lunney, Mathew S. Crowther, Mark B. Krockenberger, Mark Fisher, Linda E. Neaves

**Affiliations:** Australian Museum Research Institute, Australian Museum, 1 William Street, Sydney, New South Wales 2010, Australia; Molecular Ecology and Evolutionary Laboratory, College of Science and Engineering, James Cook University, Townsville, Queensland 4811, Australia; Centre for Sustainable Tropical Fisheries and Aquaculture, College of Science and Engineering, James Cook University, Townsville, Queensland 4811, Australia; Science for Wildlife Ltd, PO Box 5, Mount Victoria, New South Wales 2786, Australi; Centre for Tropical Bioinformatics and Molecular Biology, James Cook University, Townsville, Queensland 4811, Australia; Immunogenomics Lab, Garvan Institute of Medical Research, Darlinghurst, New South Wales 2010, Australia; JML Environmental Consultants, 16 Illallangi Close, Armidale, New South Wales 2350, Australia; School of Environmental and Rural Science, University of New England, Armidale, New South Wales 2351, Australia; Department of Planning and Environment, Locked Bag 5022, Parramatta, New South Wales 2124, Australia; School of Life and Environmental Sciences, University of Sydney, Camperdown 2006, New South Wales, Australia; Sydney School of Veterinary Science, University of Sydney, Camperdown 2006, New South Wales, Australia; 3D Ecology Mapping, Emerald Beach 2456, New South Wales, Australia; Fenner School of Environment and Society, the Australian National University, Canberra, Australian Capital Territory 2600, Australia

**Keywords:** *Phascolarctos cinereus*, phylogeography, conservation genomics, wildlife monitoring, threatened species management

## Abstract

Genetic management is a critical component of threatened species conservation. Understanding spatial patterns of genetic diversity is essential for evaluating the resilience of fragmented populations to accelerating anthropogenic threats. Nowhere is this more relevant than on the Australian continent, which is experiencing an ongoing loss of biodiversity that exceeds any other developed nation. Using a proprietary genome complexity reduction-based method (DArTSeq), we generated a data set of 3,239 high quality Single Nucleotide Polymorphisms (SNPs) to investigate spatial patterns and indices of genetic diversity in the koala (*Phascolarctos cinereus*), a highly specialised folivorous marsupial that is experiencing rapid and widespread population declines across much of its former range. Our findings demonstrate that current management divisions across the state of New South Wales (NSW) do not fully represent the distribution of genetic diversity among extant koala populations, and that care must be taken to ensure that translocation paradigms based on these frameworks do not inadvertently restrict gene flow between populations and regions that were historically interconnected. We also recommend that koala populations should be prioritised for conservation action based on the scale and severity of the threatening processes that they are currently faced with, rather than placing too much emphasis on their perceived value (e.g., as reservoirs of potentially adaptive alleles), as our data indicate that existing genetic variation in koalas is primarily partitioned amongst individual animals. As such, the extirpation of koalas from any part of their range represents a potentially critical reduction of genetic diversity for this iconic Australian species.

## 1. Introduction

The koala (*Phascolarctos cinereus*) is an iconic Australian marsupial that presents a complex management challenge because it is not uniformly threatened across its range. In 2012, the collective koala populations of Queensland (QLD), New South Wales (NSW) and the Australian Capital Territory (ACT) were classified as ‘Vulnerable’ under the Commonwealth Environment Protection and Biodiversity Conservation Act 1999 (EPBC Act) following an inquiry launched by the Australian Senate the previous year to evaluate the appropriate conservation status of the species (Senate, 2011; Shumway et al., 2015). Less than a decade later, in 2021, the status of these same populations was upgraded to ‘Endangered’ following a reassessment undertaken by the Threatened Species Scientific Committee in the wake of the unprecedented extreme fire season or “Black Summer” of 2019-2020 (TSSC, 2021). Conversely, koala populations in the states of Victoria (VIC) and South Australia (SA) are widely considered to be stable, or even overabundant in some cases, and are therefore not listed under the EPBC Act. The management of specific koala populations across Australia has been further complicated by inconsistent state-level legislative priorities and conservation planning frameworks that have, at different times, afforded the species varying levels of significance and protection (Senate, 2011; Adams-Hosking et al., 2016; McAlpine et al., 2015).

In NSW, a large body of evidence collected over nearly four decades demonstrates that koalas are experiencing widespread population declines due to a variety of threats that are often synergistic in nature (Reed et al., 1990; Lunney et al., 1995; Lunney et al., 2000; Lunney et al., 2009). Chief among these is the loss or fragmentation of critical native habitats due to land clearing, urbanisation and, more recently, extreme environmental disturbances associated with anthropogenic climate change (e.g., severe drought and altered fire regimes). Other notable threats to koalas include disease, heat waves, vehicle strikes, and dog attacks (Adams-Hosking et al., 2016; McAlpine et al., 2015, DAWE, 2022). Within this context, devising objective and unambiguous criteria for identifying and prioritising conservation targets is a crucial first step in the development of evidence-based management paradigms that will make the most efficient use of limited resources to stabilise or rehabilitate declining koala populations. To date, particular emphasis has been placed on using data-driven spatial analyses to create management divisions which represent areas and habitats with the greatest importance for the long-term persistence of the species (DECC, 2008; DPIE, 2020; DPE, 2022).

The specific criteria used to define management divisions for koalas across NSW have changed considerably over time. The ‘Recovery plan for the koala (*Phascolarctos cinereus*)’, released in 2008 by the former Department of Environment and Climate Change NSW, identified seven koala management areas (KMAs) based on a combination of local government boundaries and the known distributions of preferred koala food tree species (DECC, 2008). By contrast, the NSW Koala Strategy 2022 (DPE, 2022) does not reference these KMAs but instead identifies a total of 50 koala populations, which were derived from the 48 Areas of Regional Koala Significance (ARKS) developed by the NSW Department of Planning and Environment using state-wide information on koala occurrence (DPIE, 2020; DPE, 2022). The primary objective of defining both the KMAs and the ARKS was to create broad management areas which could be used to identify and combat threatening processes at local and regional scales. The NSW Koala Strategy 2022 further delineates the state’s koala populations into two main intervention categories. The first consists of 19 populations that are considered to be supported by sufficient information to warrant immediate targeted conservation actions. The second category covers the remaining 31 koala populations, where key knowledge gaps persist that could hinder the effectiveness of interventions to mitigate threats, enhance habitat quality, and improve overall population viability (DPE, 2022). One of the most critical of these knowledge gaps, as addressed by both the NSW Koala Strategy 2022 and the NSW Chief Scientist and Engineer’s Report, is an understanding of the mechanisms that have shaped the distribution of genome-wide genetic diversity in koalas (DPE, 2022; NSW Chief Scientist, 2016).

An extensive body of theoretical and empirical research spanning decades has established that the reduction of genetic diversity in small, fragmented populations can increase their vulnerability to extinction from both inbreeding depression and a reduced ability to adapt to rapid environmental change (Frankham, 2015; Hoffmann et al., 2020, Ralls et al., 2020). However, the importance of genetic diversity to wildlife conservation has often been neglected in both policy and practice. The NSW parliamentary inquiry (2020) into “Koala populations and habitat in New South Wales” failed to mention genetics in its 16 findings or 42 recommendations “to help ensure the future of the koala”, despite the testimony of several expert witnesses emphasising the importance of integrating genetic monitoring into ongoing management strategies (NSW parliament, 2020). Fortunately, this situation is beginning to change, with genetic and genomic approaches finding an increasingly wide range of applications in threatened species recovery efforts. These include, resolving taxonomic uncertainties (Frankham, 2010; Neaves et al., 2018; Mu et al., 2022), reconstructing historical demographic shifts (Jensen et al., 2018, Saremi et al., 2019), assessing population size and connectivity (Lowe and Allendorf, 2010; Younger et al., 2017, Hohenlohe et al. 2021), defining biologically meaningful management units (Moritz, 1994; Fraser and Bernatchez, 2002) optimising captive breeding programs (Miller et al., 2010; Witzenberger and Hochkirch, 2011), and investigating ecological or symbiotic interactions that could inform the success of future or existing conservation paradigms, such as the composition of gut microbiomes (Brice et al., 2019; Blyton et al. 2022) or host-parasite assemblages (Lott et al., 2015a; Lott et al., 2015b; Vermeulen et al., 2016a; Vermeulen et al., 2016b).

While several previous studies have investigated koala population structure and phylogeography at various scales (Houlden et al., 1999; Lee et al., 2010; Lee et al., 2011; Kjeldsen et al., 2016; Neaves et al., 2016; Dennison et al., 2017; Kjeldsen et al., 2019; Lott et al., 2022), little is currently known about the levels or distribution of genetic diversity across existing management divisions in NSW. Collecting this information is therefore not only essential for determining the overall vulnerability of regional and local koala populations to different threatening processes, and by extension their priority for targeted conservation efforts, but will also provide baseline data against which the success of future and ongoing conservation policies can be empirically assessed. The emergence of cost effective, high-throughput next-generation sequencing platforms has made the implementation of large-scale genetic monitoring programs increasingly feasible. Researchers are now able to identify thousands or even millions of hypervariable genetic markers, such as single nucleotide polymorphisms (SNPs), which can often be linked to specific regions of interest within the wider genome (Morin et al., 2004; Wright et al., 2015). Coupled with the greater availability of whole genome reference data from non-model organisms, massively parallel sequencing is facilitating the exploration of genetic diversity, population structure, and local adaptation in a wide range of threatened fauna, including koalas (Funk et al., 2012; Garner et al., 2016; Hogg et al. 2023).

In this study, we used a data set of high-quality single nucleotide polymorphisms (SNPs) generated using a reduced-representation sequencing approach to address the following three research aims: (1) analyse the fine-scale spatial genetic structure of extant populations of koalas across NSW; (2) estimate spatial patterns and rates of inter-population gene flow; (3) generate comparative genetic diversity metrics for existing management divisions (ARKS), and test for relationships between the quality of koala habitat and overall levels of genetic diversity.

## 2. Methods

### 2.1 Sample Collection

Blood, tissue and buccal swab samples, representing 314 individuals from 29 of the 48 ARKS (corresponding to 14 of the 19 populations for immediate investment, and 16 of the 31 populations with key knowledge gaps), were obtained from researchers, environmental consultants, veterinarians, and wildlife rehabilitators from across the state of NSW (Table S1.1; Figure 1). This constitutes the most geographically comprehensive survey of genetic diversity in NSW koalas to date. Where possible, sampling gaps were also filled by sourcing archived biological material from the Australian Museum Koala Tissue Biobank, the designated repository for koala tissue and genetic material obtained in NSW. All samples were stored in 70%–100% ethanol or frozen at −80°C prior to DNA extraction and genotyping.

**Figure 1.**
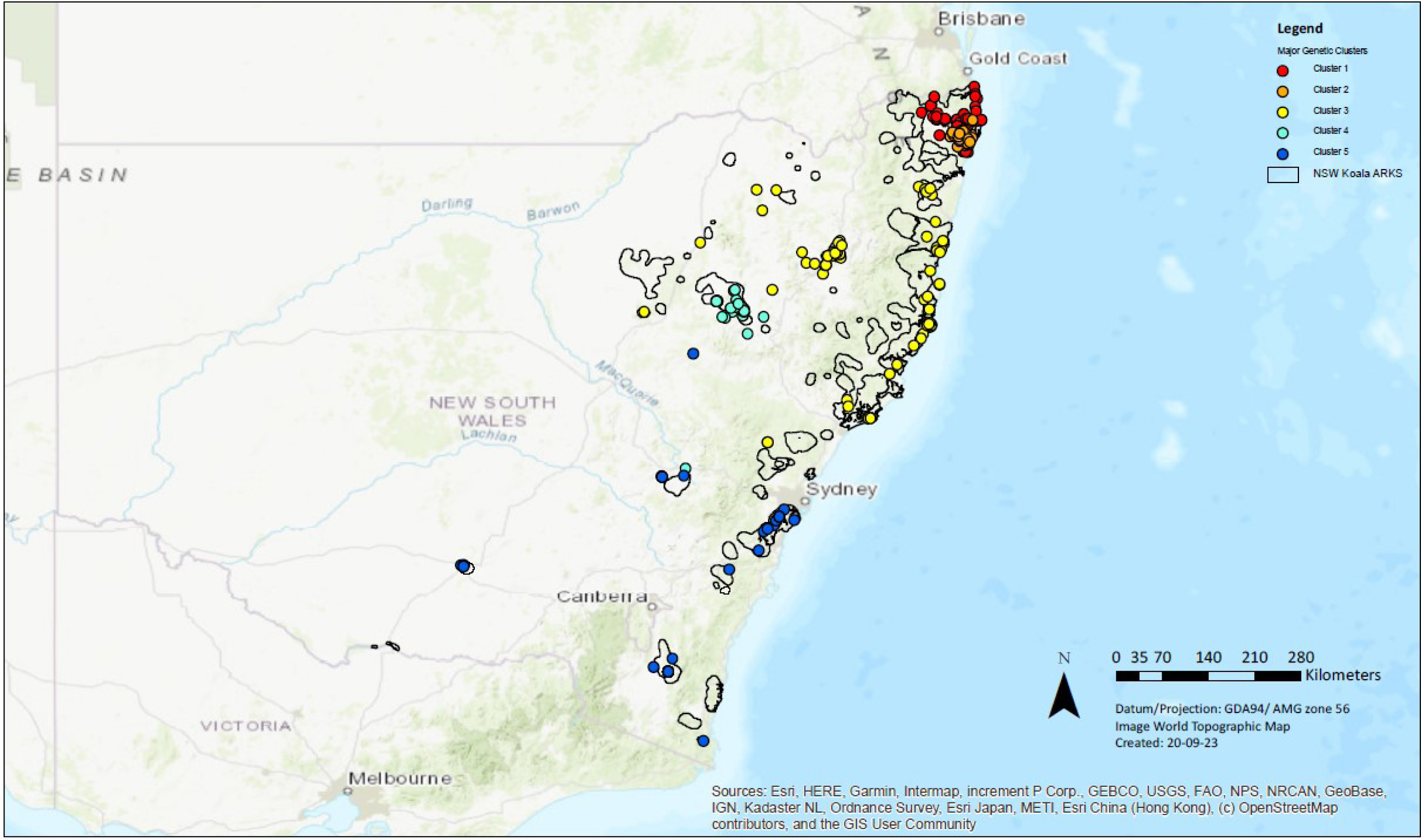
Distribution of the 314 koala specimens included in this study relative to the 48 Areas of Regional Koala Significance (ARKS). Samples are colour coded by major genetic cluster of origin, as identified by both DAPC and STRUCTURE.

### 2.2 DNA extraction and genotyping

Genomic DNA was extracted using either the Bioline Isolate II Genomic DNA Kit (Bioline, Eveleigh, Australia) following the manufacturer’s protocols, or a standard high-salt precipitation procedure (Sunnucks and Hales, 1996). Genotyping was performed using the Diversity Arrays Technology platform (DArTseq™). DArTseq is a restriction enzyme-based genome complexity reduction method that has been utilised to generate SNP data in a wide range of vertebrate species for phylogeographic, phylogenetic, and population genetic studies. DNA was processed as per Kilian et al. (2012), using paired adaptors which corresponded to two different restriction enzyme overhangs: PstI and SphI. The PstI-compatible adapter included an Illumina flow cell attachment sequence, a sequencing primer binding site, and a varying length barcode region. The reverse adapter contained a SphI-compatible overhang sequence and a flow cell attachment region. A digestion–ligation reaction was performed at 37 °C for 2 h with ∼100–200 ng of gDNA per sample. The DNA fragments that were successfully cut by both PstI and SphI were then amplified by 30 cycles of polymerase chain reaction (PCR), and the PCR products were sequenced as 77-bp or 138-bp single-end reads on the HiSeq 2500 and Novaseq 6000 platforms, respectively (Illumina, San Diego, USA). After demultiplexing and adapter trimming, the short-read sequence data were processed using Stacks v2.64. Sequencing reads were standardised by truncating them to 69bp in length and low-quality data (based on the PHRED scores provided in the FASTQ files) were identified and discarded using the process_radtags program (Catchen et al. 2013). Sequencing reads were discarded when the probability of them being correct dropped below 99.9% (i.e., a PHRED score of 30). Prior to implementing ref_map.pl in Stacks, the cleaned FASTQ files from the previous step were aligned to the koala reference genome (GCA_002099425.1_phaCin_unsw_v4.1, Johnson et al. 2018) using the mem function in Burrows-Wheeler Aligner (BWA) v0.7.15 (Li and Durbin, 2010; Willet et al., 2021). These alignments were subsequently converted to BAM format using SAMtools v1.6 (Li et al., 2009). The reference-aligned data were then used to assemble the sequences into loci and identify SNPs using the *ref_map.pl* pipeline in Stacks (Catchen et al. 2013). Briefly, this pipeline aligns matching sequences into ‘stacks’, which are in turn merged to form putative loci. At each of these loci, nucleotide positions are examined, and SNPs are called using a maximum likelihood framework. A catalogue is then created of all possible loci and alleles, against which the individual samples are matched. The *ref_map.pl* pipeline was implemented using the default parameters, with one exception: the alpha threshold required to call a SNP was reduced from 0.05 to 0.01 (i.e., a greater number of sequence reads were required to make a SNP call statistically significant at each locus) in order to minimise the risk of introducing markers that represented false positives into the data set. Similarly, to reduce the probability of linkage between markers, a single SNP was extracted from each locus using the *populations* program in Stacks. The entire procedure, from library preparation to SNP calling, was repeated a second time for 60 technical replicates. Only biallelic loci with 100% reproducibility (i.e., no genotyping errors) were retained. Further filtering of the genotypes was then performed in PLINK v 1.9 (Purcell et al., 2007). Variant sites with call rates of <90% and minor allele frequencies of <0.005 were removed from the data set. This MAF threshold was chosen to reduce the probability of including false alleles originating from sequencing error by guaranteeing that each allele was sampled in in ≥2 individuals independently (as shown by the formula 3/2 N: 3/(2 × 317) = 0.005) (Wright et al., 2019; Lott et al., 2020). Finally, to accommodate downstream genetic analyses requiring a neutral set of markers, the data set was further filtered to remove SNPs out of Hardy-Weinberg equilibrium, and outlier SNPs that potentially represented loci under selection. Candidate outlier SNPs were identified using the program PCAdapt for the R-software (Luu et al., 2017). PCAdapt employs a Bayesian hierarchical factor model to describe population structure as latent factors, and locus-specific effects on population structure as correlated factor loadings (Duforet-Frebourg et al., 2014). Unlike many alternative outlier detection models, PCAdapt bypasses the assumption of an island model of gene flow and avoids the need to define population structure a priori. Based on scree plots depicting the proportion of explained variance (Figure S2.1), 4 K populations were chosen to account for neutral structure in the data set using Cattell’s rule, which states that components corresponding to eigenvalues to the left of the straight line should be kept (Cattell 1966). Outlier loci were scored based on Bonferroni corrected p-values and a stringent false detection rate threshold of 0.01 was selected. After controlling for the effects of neutral population structure, a total of 13 loci were identified as candidates for being under selection and removed from the data set. Departure from Hardy–Weinberg equilibrium (HWE) was tested for each locus using the package pegas v0.12. for the R software (Paradis, 2010). However, as the failure to consider existing population structure by HWE filters has been shown to result in heterozygote deficiencies at potentially informative loci due to Wahlund effects (De Meeûs, 2018), genetic structure was first assessed by performing a discriminate analysis of principal components (DAPC). The primary advantage of DAPC is that it does not rely on a particular population genetics model and is therefore free from assumptions about HWE or linkage disequilibrium. The major genetic clusters identified by DAPC were then used as the basis for partitioning samples for HWE filtering. While sampling locations are more commonly used as a proxy for genetic populations in the literature, such an approach might artificially inflate divergence estimates between sampling locations if they do not accurately reflect the underlying population structure (Pearman et al. 2022). Consequently, to maximise the retention of potentially informative loci, while also accommodating downstream analyses which require neutral genetic markers, we elected to only remove loci that deviated from HWE in all major genetic clusters identified by DAPC. The quality control criteria described above resulted in a data set of 3,239 high quality SNPs that, except where specifically indicated, were used for all downstream analyses.

### 2.3 Fixed difference analysis

To examine the possibility that some existing management divisions might represent demographically independent units characterised by restricted gene flow, a fixed difference analysis was performed in dartR (Gruber et al., 2018) using the default parameters. A fixed difference occurs when two populations share no alleles at a particular locus. Therefore, the accumulation of fixed differences between populations strongly indicates a lack of gene flow. We elected to partition koala samples by ARKS for this and all other analyses requiring a priori assignment of individuals into specific management divisions, as the criteria that were used to develop them are transparent and well documented. Conversely, the rationale for modifying the 48 ARKS into the 50 populations outlined in the NSW Koala Strategy has not been published. Using the *gl.collapse.recursive* function (Gruber et al., 2018), fixed differences were summed over pairwise groupings of populations (i.e., ARKS). When no fixed differences were detected between the two populations in question, they were amalgamated. This process was repeated until no further consolidation was possible. As noted by Georges et al. (2018), the decision to amalgamate two populations can be made with relative certainty, but the separation of two populations based on the detection of one or more fixed differences can be influenced by false positives that may arise as a consequence of the finite sample sizes involved. As such, the groupings of ARKS identified by the fixed-difference analysis described above were tested for significance. Population pairs for which the number of fixed differences was not statistically significant (i.e., the observed number of fixed differences was not significantly different from the expected rate of false positives) were further amalgamated. The fixed difference analysis was performed using a data set that retained SNPs that were putatively under selection and/or out of HWE (i.e., unfiltered), as removing these loci could potentially inflate the counts of fixed allelic differences between ARKS.

### 2.4 Analysis of population structure

The fine-scale population structure and admixture history of koalas across NSW was investigated using a Bayesian model-based clustering approach implemented in the program STRUCTURE 2.3.4 (Pritchard et al., 2000). Ten independent runs were used to model up to 15 populations (i.e., K = 1–15), with each run consisting of a burn-in period of 10^5^ iterations, followed by 2 × 10^5^ Markov chain Monte Carlo (MCMC) replicates. We did not use location information to establish priors and the chosen ancestry model assumed both admixture and correlated allele frequencies. Structure Harvester (Earl and Von Holdt, 2012) was then used to determine the optimal value of K by calculating both the maximum delta log likelihood (Δ*K* Evanno et al., 2005) and the maximum posterior probability (*L*(*K*) Pritchard et al., 2000). Finally, the Cluster Markov Packager Across K (CLUMPAK) web server (Kopelman et al., 2015) was used to merge and visualise replicate runs as bar plots. Following the recommendations of Janes et al. (2017), each of the clusters identified by this procedure was subsequently rerun to test for additional sub-structuring. For all downstream analyses, individual samples were categorised into distinct populations according to their membership coefficients, defined here as the proportion of each genotype that could be attributed to a particular genetic cluster. Admixed koalas were assigned to the population which accounted for the largest percentage of their genome.

To quantify interpopulation genetic similarity, pairwise *F_ST_* indices and their 95% confidence intervals were calculated with 1000 bootstraps in the R package dartR (Gruber et al., 2018). The hierarchical partitioning of genetic variation was assessed using an analysis of molecular variance (AMOVA) in the R package Poppr version 2.7.1 (Kamvar et al., 2014). Finally, the correlation between geographical and genetic distances was examined using a Mantel test performed in dartR (Gruber et al., 2018).

### 2.5 Estimating contemporary inter-population gene flow

Contemporary migration patterns were further investigated in BayesAss version 3.0 (Wilson and Rannala, 2003). This software program utilises a Bayesian statistical framework to estimate recent immigration rates from multilocus genotypes. Following the recommendations of Meirmans, (2013), sampling locations were pooled (i.e., fewer populations with many individuals) to increase the statistical power of the analyses. The koala samples were partitioned into a total of six pools, with four of these pools directly corresponding to a major genetic cluster identified by DAPC and STRUCTURE. The final two pools were created by subdividing the fifth major genetic cluster into two groups in order to separate samples sourced from collection sites to the east and west of the Great Dividing Range (GDR; Table S1.1). This was done to test the hypothesis of Lott et al, (2022) that a source-sink population dynamic currently exists across the GDR. To achieve the recommended acceptance rates (0.2-0.4), the mixing parameters for the inbreeding coefficient and allele frequency were set to 0.10 and 0.30 respectively. Analyses were then run for 2 × 10^7^ iterations, with a burn-in period of 5 × 10^6^ iterations, and a sampling frequency of 2,000. The analysis was repeated five times with different starting-seed values. Convergence was diagnosed using two different approaches: first, by confirming that mean parameter estimates were consistent between replicate runs and, second, by ensuring that large log-probability fluctuations were confined to the burn-in phase and that no major oscillations occurred which might influence parameter estimates (Figure S2.2). The median migration rates of the five independent BayesAss runs were used to construct 95% credible sets by multiplying the mean standard deviation for each migration rate by 1.96, as suggested in the BayesAss user manual. Migration rates were considered to be significant when the credible set did not overlap with zero.

### 2.6 Comparative genetic diversity metrics

Homozygosity by locus (HL), an individual-based measure of genetic variation, was calculated for each sample using the GENHET function (Coulon, 2010) in R. The primary advantage of this method is that it accounts for allelic variability when weighing the contribution of each locus to the homozygosity index (i.e., greater weight is given to the most informative loci) (Aparicio et al., 2006). Consequently, HL is expected to be more strongly correlated with inbreeding coefficients and genome-wide homozygosity than other commonly used individual-based measures of genetic variation such as the uncorrected proportion of homozygous loci or internal relatedness (IR) (Aparicio et al., 2006), particularly in open populations with varying levels of dispersal and/or admixture, as is likely to be the case in koalas. Mean HL values were calculated for both the 29 sampled ARKS, and the five major genetic clusters identified by DAPC and STRUCTURE.

To investigate the relationship between genetic diversity in koalas and several key aspects of their environment we employed a multilevel mixed-effects linear model, where HL was modelled as the response variable (R Core Team, 2014). Information on the basic characteristics for each ARK was sourced from the Koala Habitat Information Base (DPIE, 2019; Table S1.2). Three key predictors were included in the final model: the percentage of high and moderate functional habitat, the percentage of low and very low functional habitat, and human population density. Additionally, as the effects of these predictors are unlikely to be entirely independent, we included several interaction terms in our final model, specifically: the percentage of high and moderate functional habitat and the percentage of low and very low functional habitat, the percentage of high and moderate functional habitat and human population density, and the percentage of low and very low functional habitat and human population density. These interaction terms appear as bilinear functions of the paired predictors. Finally, the major genetic clusters from which individual koala samples were sourced were fitted as random effects. Prior to analysis, the continuous data were scaled to a range between 0 and 1, while the categorical variable was recoded into a set of separate binary variables (i.e., dummy coding). The model outputs were compared using an analysis of deviance table.

## 3. Results

With the exception of Wollemi National Park, all of the sampled ARKS collapsed into a single operational taxonomic unit (OTU) based on corroborated fixed differences. While the two identified OTUs differed by only two fixed differences, subsequent testing supported their significance (false positive expectation = 0.4, p < 0.001). It is important to note, however, that the false positive rate for fixed differences is largely a product of sample size. As the Wollemi National Park ARKS was represented by only two specimens, these results must be interpreted extremely cautiously as it is highly probable that they represent a false positive.

### 3.2 Population structure

The initial DAPC analyses supported the existence of five major genetic clusters in koalas across the state of NSW (Figure 2). This was consistent with the results of the Bayesian model-based clustering procedure implemented in STRUCTURE which indicated that either two (maximum Δ*K*) or five (maximum *L*(*K*)) populations best described the distribution of genetic variability (Figure 3). Furthermore, re-running STRUCTURE separately for each of the sample groups identified by the K=5 solution uncovered an additional putative sub-cluster. Given that the demographic, environmental and historical processes that have led to the current distribution of genetic diversity in koalas are likely to be multifaceted and complex, it is perhaps unsurprising that different levels of organisation would be present in the genetic structure. Determining the clustering solution which best describes population structure in koalas is therefore a non-trivial task. Janes et al. (2017) has shown that Δ*K* tends to underestimate population structure by identifying only the highest level of differentiation. Conversely, Perez et al. (2018) demonstrated that STRUCTURE outputs can be heavily influenced by isolation-by-distance, most often through the detection of artificial and misleading genetic clusters. However, given that the STRUCTURE and DAPC analyses both converged on five clusters as best describing the distribution of genetic diversity in contemporary NSW koala populations, a value of five was used for any downstream analyses that incorporated assumptions of population structure.

**Figure 2.**
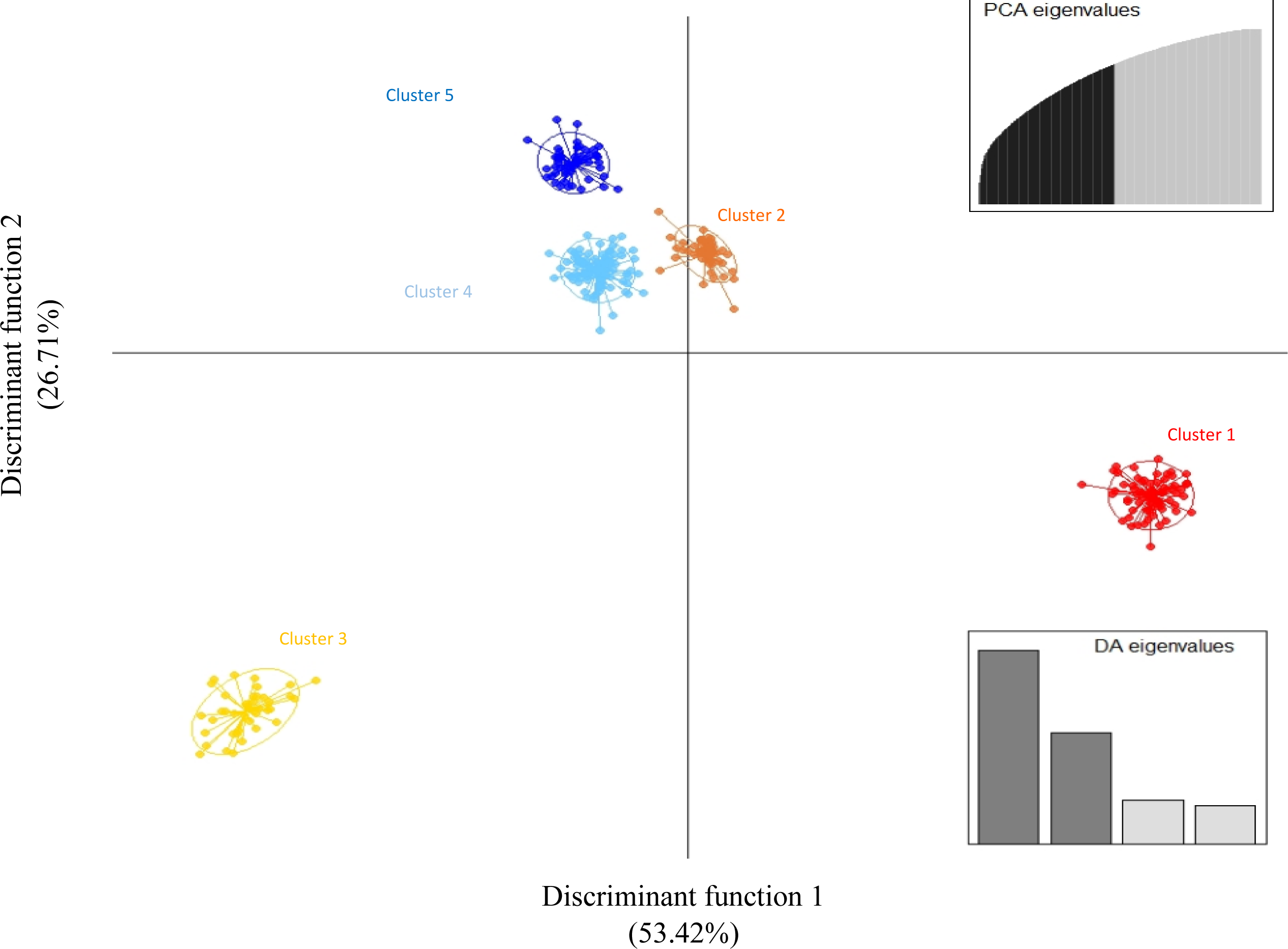
Scatterplot showing the first two principal components of the discriminant analysis of principal components (DAPC) which was applied to the data set prior to filtering the loci that were out of Hardy-Weinberg equilibrium. Each koala genome is represented by a single dot, while inertial ellipses are depicted as ovals, with the lines extending to the centroids of each cluster. The number of principal components retained for each analysis (PCA) and the relative amount of genetic variation contained in each discriminant factor (DA) are shown using eigenvalue plots.

**Figure 3.**
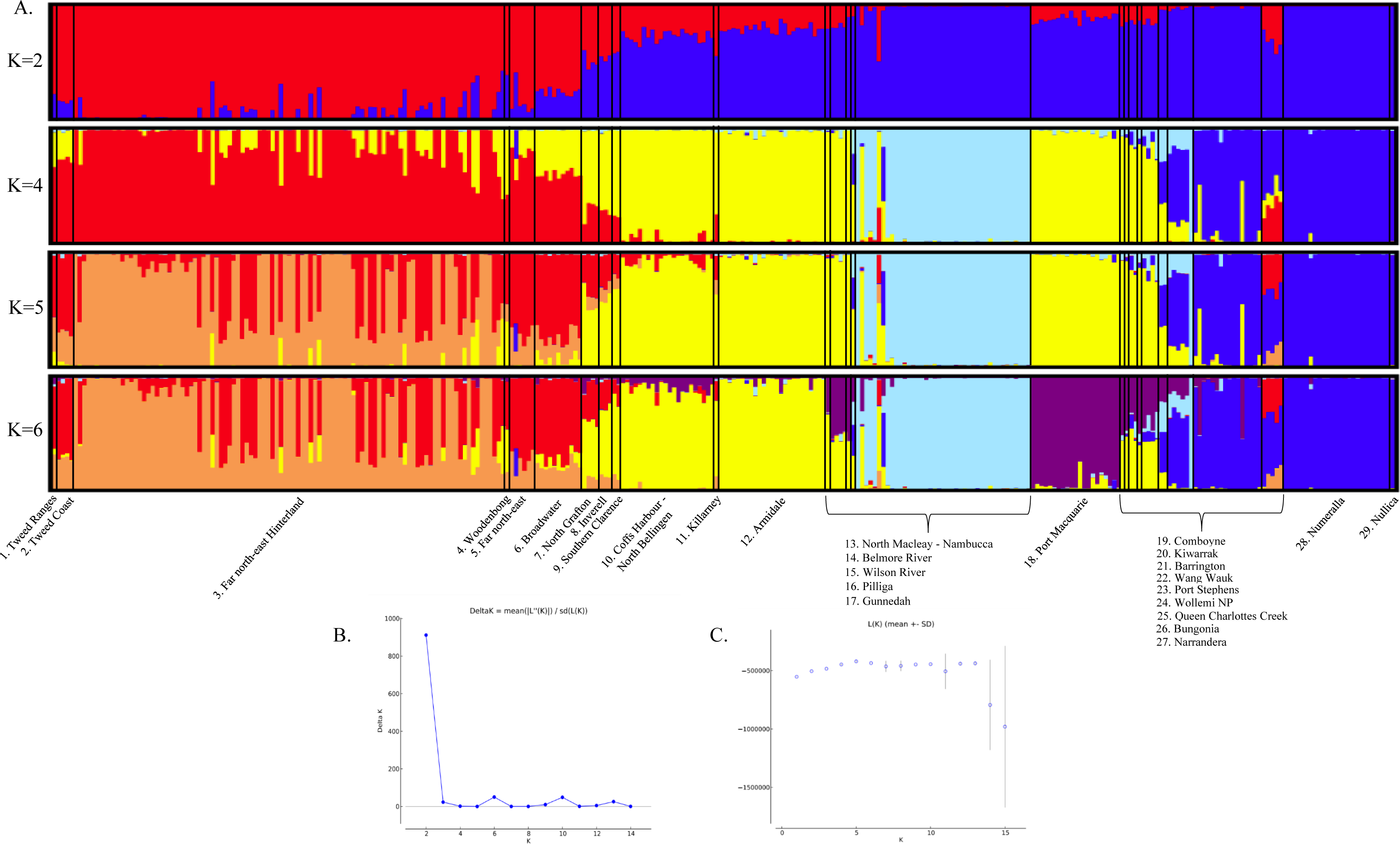
Distribution of genetic diversity in contemporary koala populations across the state of NSW. A) STRUCTURE plots showing the inferred ancestry proportions for 314 koalas sourced from 29 Areas of Regional Koala Significance (ARKS) at four different values of K. Optimal clustering solutions according to B) maximum delta log likelihood (Δ*K*) and C) maximum posterior probability *L*(*K*).

The pairwise *F*_ST_ values indicated low to moderate differentiation between the five major genetic clusters of koalas (Table 1). The AMOVA demonstrated that the greatest proportion of genetic variance occurred within individual koalas (69.280%), while variance between individuals within the 29 ARKS and between ARKS within the five major genetic clusters accounted for 8.977% and 11.918% of genetic diversity, respectively (Table 2). In contrast, genetic structure among the five major population clusters represented 9.825% of total variance. Finally, the Mantel test indicated that a positive correlation existed between genetic distances and geographic distances in NSW koalas (r = 0.573, p = 0.025).

**Table 1.** Pairwise genetic differentiation (*F*_ST_) between the five major genetic clusters of koalas (bottom-left diagonal) and their associated Bonferroni-corrected *p*-values (top-right diagonal).

|  | Cluster 1 | Cluster 2 | Cluster 3 | Cluster 4 | Cluster 5 |
| --- | --- | --- | --- | --- | --- |
| Cluster 1 | – | 0.073-0.085 | 0.082-0.095 | 0.198-0.222 | 0.173-0.194 |
| Cluster 2 | 0.079 | – | 0.156-0.177 | 0.277-0.307 | 0.240-0.266 |
| Cluster 3 | 0.088 | 0.166 | – | 0.134-0.153 | 0.120-0.136 |
| Cluster 4 | 0.21 | 0.292 | 0.143 | – | 0.183-0.206 |
| Cluster 5 | 0.183 | 0.253 | 0.128 | 0.194 | – |

**Table 2.**
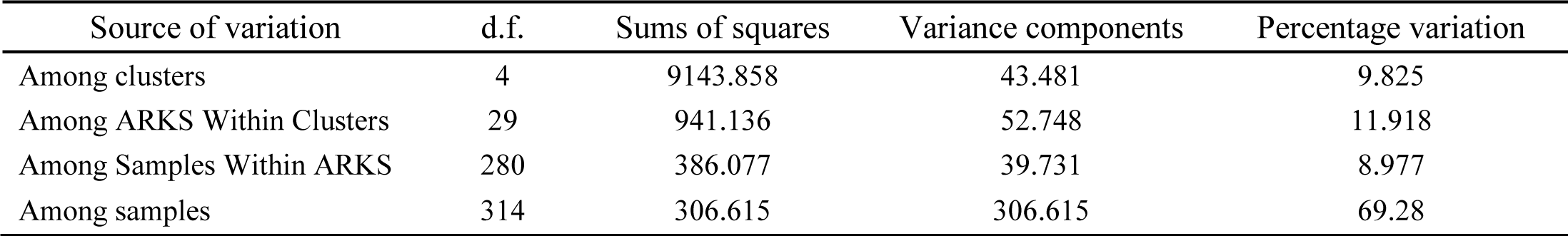
Hierarchical AMOVA results showing levels of genetic structure among the five major genetic clusters identified by STRUCTURE/DAPC, the areas of regional koala significance (ARKS), and individual animals (n=314).

### 3.3 Estimating inter-population gene flow

Bayesian estimations of contemporary migration rates between the major genetic clusters of koalas varied but were generally low and highly asymmetrical (Figure 4; Table S3.1). The largest proportion of migrants appeared to be the result of unidirectional dispersal from Cluster 1 into Cluster 2 (0.250 ± 0.032). This is consistent with the high levels of admixture that were observed between these two major genetic clusters. Low but statistically significant northward dispersal was also detected between Cluster 3 (East GDR) and Cluster 1 (0.058 ± 0.028). Additionally, Cluster 3 (East GDR) was found to be a significant source of migrants for the western koala populations that constituted Cluster 3 (West GDR) (0.208 ± 0.039). The movement of koalas appeared to be highly asymmetric, as comparable levels of southward and eastward dispersal were not detected. The 95% credibility intervals of migration rates between all other major genetic clusters encompassed zero and were therefore interpreted to be nonsignificant.

**Figure 4.**
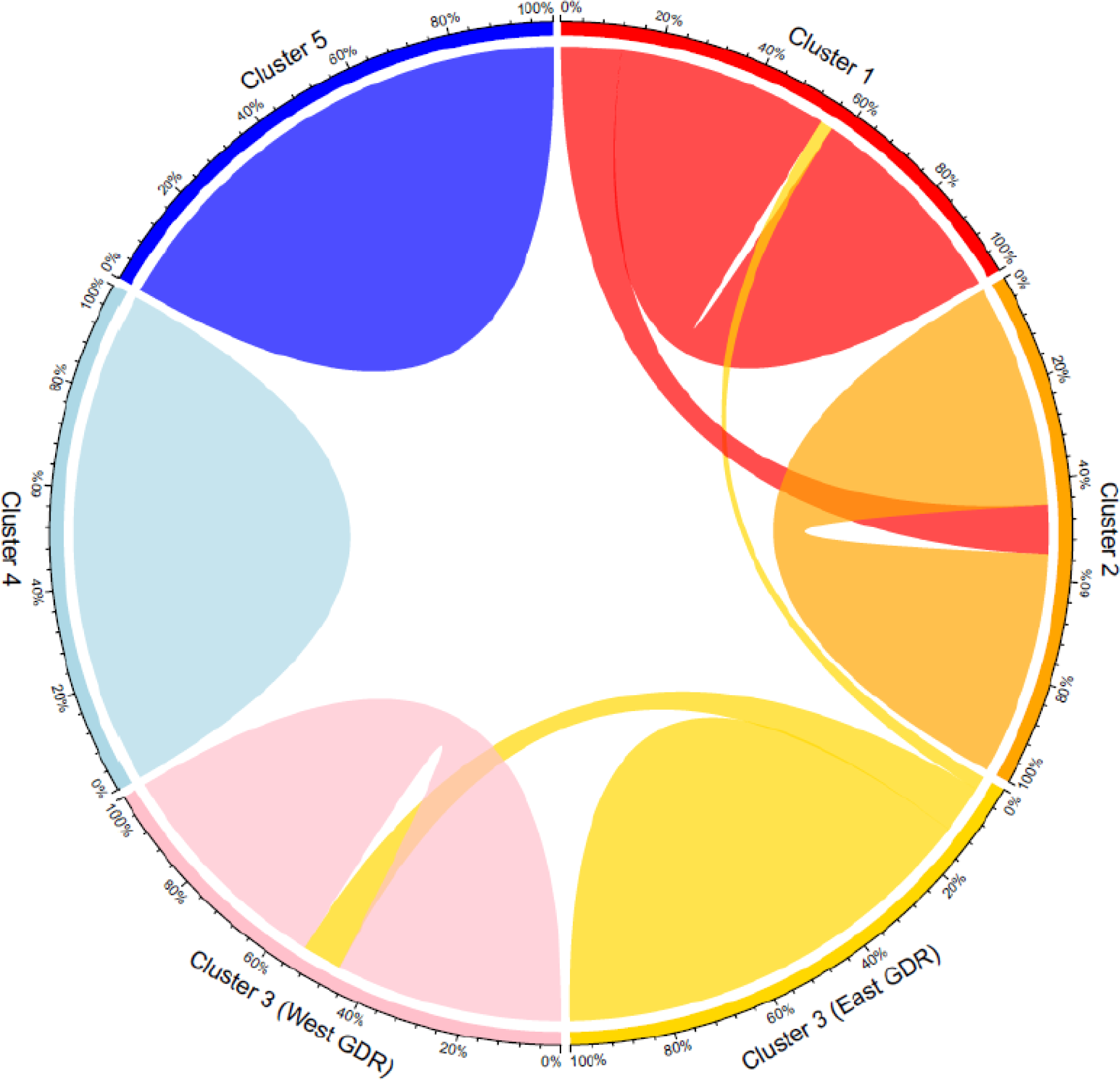
Chord diagram depicting contemporary gene flow estimates for NSW koalas derived from BayesAss (raw BayesAss results are displayed in Table S3.1 in Supplementary Material). The thickness of the chords represents the rate of migration from the source to the recipient population/s; only migration rates significantly different from zero are displayed. The proportion of migrants relative to total population size is depicted on the outer section axis.

### 3.4 Comparative genetic diversity metrics

Homozygosity by locus varied across the 29 sampled ARKS (Figure 5). While precise estimates of genetic diversity should be interpreted cautiously when sample sizes are small, the Gunnedah, Port Stephens, Queen Charlottes Creek, and Numeralla ARKS all exhibited HL values that were significantly higher than the state average. Furthermore, a geographical pattern of genetic diversity emerged in which HL values increased with latitude. Of the five major genetic cluster identified by the DAPC and STRUCTURE analyses, Cluster 4 (centred on Gunnedah and the Liverpool Plains) had the lowest overall genetic diversity, while Cluster 1 (in far north-east NSW) had the highest. Multilevel mixed-effects linear models indicated that, when major genetic cluster of origin was controlled for, neither the chosen predictors nor their interaction terms were significantly correlated with genetic diversity in koalas (Table 3).

**Figure 5.**
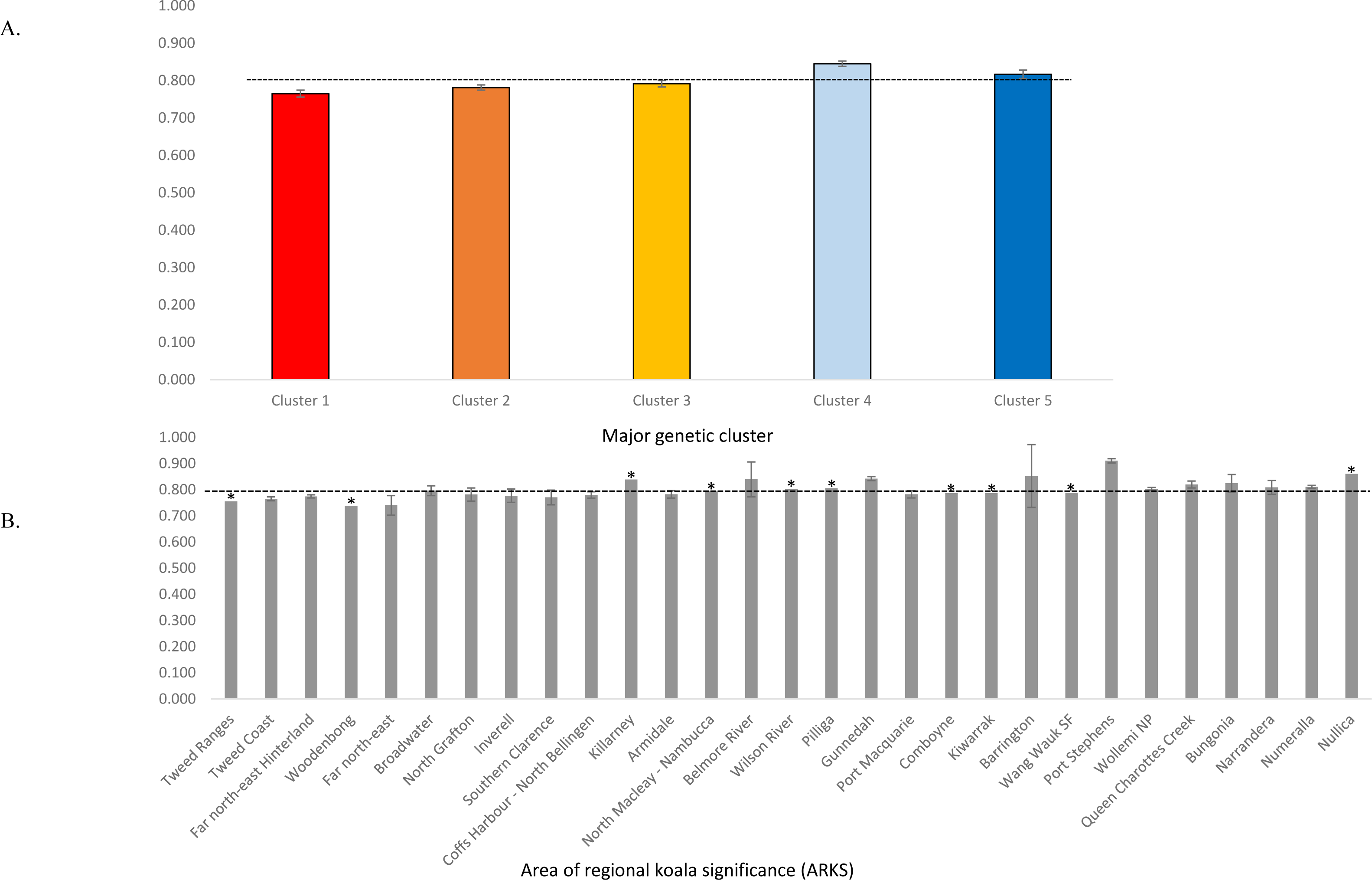
Mean homozygosity by locus (HL) scores and associated 95% confidence intervals for A) the five major genetic clusters identified by DAPC/STRUCTURE and B) 29 Areas of Regional Koala Significance (ARKS). In both instances, the mean state-wide HL is represented by a broken black line. ARKS represented by a single sample are denoted with an asterisk, indicating that these values cannot be taken as representative of the overall levels of genetic diversity within these management divisions.

**Table 3.**
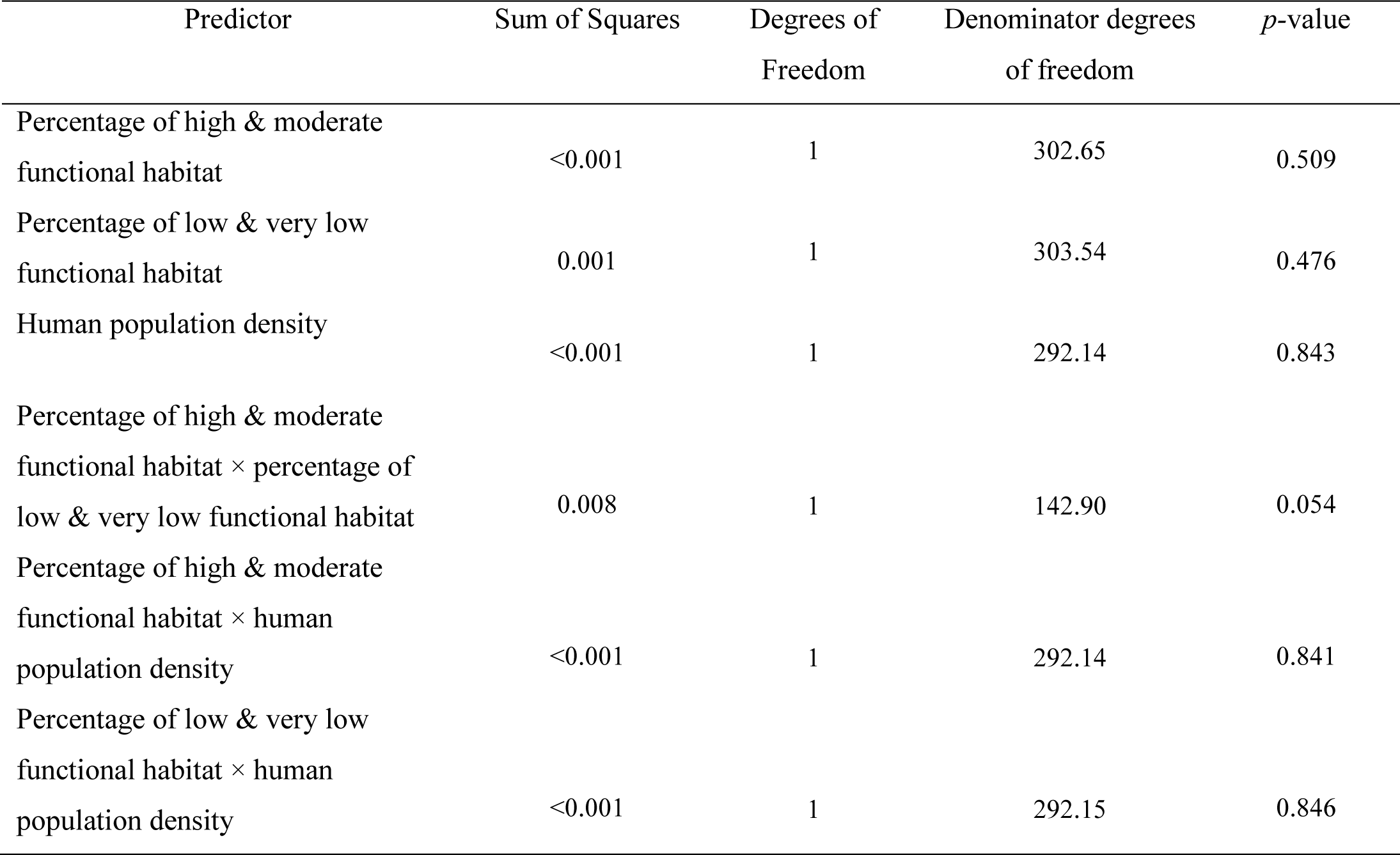
Analysis of variance using Satterthwaite’s method for changes in homozygosity by locus (HL) based on multilevel mixed-effects linear modelling.

## 4. Discussion

### 4.1 Koala population structure in NSW

Significant genetic structuring was detected in koala populations across the state of NSW. Both multivariate and model-based clustering analyses indicated that there are five extant major genetic clusters, with additional hierarchical structure identified within at least one of these clusters. This distribution of genetic diversity broadly corresponds to that described previously by Kjeldsen et al., (2016), Johnson et al., (2018) and Lott et al., (2022). Perhaps unsurprisingly, vicariance across biogeographic barriers for forest adapted taxa appears to have been an important driver of genetic differentiation in NSW koala populations. A prominent north-south separation was observed across the Sydney Basin region, with the southernmost lineage (Cluster 5) corresponding to what has been termed the South Coast NSW cluster (Johnson et al., 2018; Kjeldsen et al., 2019; Lott et al., 2022). The Sydney Basin and neighbouring Illawarra region are defined by extensive, low-lying coastal plains which are boarded to the west by both the Blue Mountains and a region of uplifted sandstone known as the Illawarra Escarpment (Bryant and Krosch, 2016). It has been speculated that the vicariance of coastal forest habitat across these areas during the Miocene-Pleistocene may have played an important role in structuring genetic diversity within a wide array of vertebrate species, including koalas (Sumner et al., 2010; Pepper et al., 2014; Frankham et al., 2016).

Further north, three distinct genetic clusters of koalas were found to inhabit coastal NSW. The distribution of one of these lineages (Cluster 3) appeared to match that of the Mid-Coast NSW genetic cluster previously identified by Johnson et al., (2018) and Lott et al., (2022), which is hypothesised to be bordered by the Clarence River Corridor in the north, and either the Hunter Valley or the Sydney Basin in the south. Both of these putative biogeographic barriers are lower elevation zones of dry, warm, open woodland or grassland that are expected to represent a significant obstacle to the movement of koalas. Cluster 3 was also found to extend west of the GDR, where an additional koala lineage (Cluster 4) was identified, which was apparently centred on the Liverpool Plains in the North-West Slopes region of NSW. There are several possible reasons why two distinct genetic clusters may have been detected across these regions where Johnson et al., (2018) and Lott et al., (2022) previously reported only one. The first is that the intensive, local-level sample collection paradigm employed in this study may have allowed the detection of additional population structure across the state of NSW that was masked from previous, continent-wide genetic surveys of koalas. Alternatively, it may be because the aforementioned genetic studies were based on the analysis of SNPs derived exclusively from exons (protein coding gene regions). The level of genetic diversity found in exons is often lower than in non-protein coding regions of the genome (e.g., introns). This has variously been attributed to stronger purifying selection, higher mismatch repair activity or some combination of the two (Frigola et al., 2017). In practice, this means that exons in biogeographically isolated koala populations can be expected to retain their identity longer than non-protein coding regions, and analyses based on molecular markers derived from these parts of the genome may therefore be biased towards the detection of older divisions between populations. Conversely, molecular markers primarily derived from non-protein coding regions (as, statistically, is likely to be the case with the DArTSeq data set used in this study) may reflect population structure that has arisen comparatively more recently, possibly even as a consequence of anthropogenic habitat fragmentation.

The final two NSW koala lineages identified by our analyses (Cluster 1 and Cluster 2), which were coincident with the South-East Queensland genetic cluster previously identified by Johnson et al., (2018) and Lott et al., (2022), did not appear to be strongly associated with vicariance across any of the biogeographic barriers which have been proposed to exist in this region. While it is unclear at present how these two lineages have maintained their unique genetic identities despite their close geographical proximity, one possibility is that it is a consequence of historically restricted dispersal caused by the expansion of rainforest habitat in the vicinity of the McPherson and Border Ranges during the Pleistocene interglacials (Bryant and Krosch, 2016; Flores-Rentería et al., 2021). Alternatively, higher-than-average recruitment in natal colonies and other social factors may be reinforcing philopatry-related genetic structure. However, further research would be required to either confirm or disprove this interpretation.

### 4.2 Spatial patterns of gene flow

Mantel tests revealed that a significant relationship exists between genetic distances (*F_ST_*) and geographical distances in koalas. Genetic isolation-by-distance is therefore likely to be a significant driver of regional variation in NSW koala populations. However, the detection of genetic structuring across relatively small areas also suggests that koala dispersal can be impeded by features of the landscape. These two scenarios are by no means mutually exclusive, and it seems clear that the movement of individuals between vicariant genetic clusters, however infrequently, has resulted in complex patterns of genetic diversity across the state of NSW. Comparatively rare, long-distance dispersal events, possibly coupled with ancestral range expansions from the isolated refugia that are hypothesised to have existed during one or more of the Pleistocene glacial periods (Adams-Hosking et al., 2011; Lott et al., 2022), appear to have resulted in a state-wide isolation-by-distance effect which reflects the once continuous geographic distribution of koalas across NSW. However, this isolation-by-distance effect has failed to obscure the pre-existing genetic structure caused by vicariance across more ancient biogeographic barriers.

Despite evidence for widespread admixture between the major genetic clusters, contemporary gene flow was generally limited. The koala populations of the Liverpool Plains (Cluster 4) and to the south of the Sydney Basin (Cluster 5) were found to be particularly isolated, with no evidence for effective dispersal in recent generations. Koalas to the west of the GDR are characterised by an increasingly disjunct and scattered distribution, and our findings would be consistent with habitat fragmentation limiting gene flow between previously interconnected regions and populations. Conversely, there is reason to believe that koalas in both these areas (i.e., the Liverpool Plains and southern NSW) have long been relatively isolated from the rest of the state. It has been hypothesised that the more marginal habitat towards the western edge of the koala’s distribution has historically supported low density, widely dispersed populations that have only transiently increased in size following periods where climatic conditions (e.g., rainfall) have briefly improved (Ellis et al., 2017; Lunney et al., 2012; Lunney et al., 2017; Predavec et al., 2018; Lunney et al., 2020). The absence of large, stable patches of functional habitat to support long-term population growth and provide corridors for effective dispersal may have served to reinforce the genetic distinctiveness of western edge koala populations relative to their coastal conspecifics. Similarly, the apparent isolation of koala populations in southern NSW may indicate that the heterogenous landscapes of the Sydney Basin/Illawarra region have historically represented a greater obstacle to dispersal than the biogeographic barriers that have been hypothesised to exist in other parts of the state. It is also highly likely that additional barriers to gene flow have been created by the widespread urbanisation and land clearing that has occurred across these areas since the European colonisation of Australia (Lunney and Leary, 1988; Lunney et al., 2010; Lunney et al., 2014). While the koala populations of the Liverpool Plains and to the south of the Sydney Basin/Illawarra region appear to have been effectively isolated from the rest of the state in recent generations, the underlying barriers to dispersal have clearly not always been absolute. Small numbers of highly admixed individuals were detected within both groups, indicating that they may have once enjoyed greater, although perhaps still limited, connectivity to genetically distinct koala populations across NSW. Alternatively, undocumented translocations could have introduced some genotypes into areas where they would otherwise not be expected to occur.

While contemporary gene flow between the major genetic clusters was generally low or absent, there were several notable exceptions. Koala populations from the Mid North Coast (Cluster 3) were found to be a significant source of migrants to multiple neighbouring regions, including those to the west of the GDR. While the precise mechanisms underlying state-wide dispersal patterns remain unclear, these observations strongly suggest that a source–sink dynamic exists in this species, whereby the relatively large and stable koala populations occupying high-quality coastal habitats are contributing a disproportionate number of immigrants to less densely populated regions in the west. Identifying populations or major genetic clusters that are net exporters of immigrants has important implications for koala conservation. Extreme weather events, such as drought and heat waves, are strongly associated with poor health and increased mortality in koalas, particularly for populations living near the arid edge of the species’ current distribution (Adams-Hosking, et al., 2011; Lunney et al., 2012; Davies et al., 2013; Davies et al., 2014; Seabrook et al., 2014; Lunney et al., 2014; Lunney et al., 2020). If the koalas occupying marginal habitats are disproportionately vulnerable to periodic population crashes caused by long-term fluctuations in temperature and rainfall, then the asymmetric dispersal of individuals from larger, self-supporting coastal populations may help facilitate their recovery by maintaining genetic diversity and overall population viability in subsequent generations. Consequently, failure to conserve koala source populations, and the critical native habitats that supports them, may also negatively affect the survival of dependent sinks towards the western edge of koala distribution.

### 4.3 Genetic diversity metrics

With several exceptions, genome-wide genetic diversity did not differ significantly between the sampled ARKS. Notably, the Gunnedah, Port Stephens, Queen Charlottes Creek, and Numeralla ARKS all exhibited levels of genetic diversity that were significantly lower than the state average. While further sampling is required to confirm some of these findings, this may indicate that koalas in these ARKS are more vulnerable to key threatening processes than populations in other regions. When examining the five major genetic clusters, the western (Cluster 4) and southern (Cluster 5) most lineages were found to have the lowest overall levels of genome-wide genetic diversity. These results largely support the findings of Johnson et al., (2018) and Lott et al., (2022) which demonstrated that, on a continental-scale, koala genetic diversity decreased along a north-south cline. In the past, this phenomenon has been attributed primarily to hunting or habitat-loss associated population bottlenecks following European colonisation of Australia. However, mounting evidence suggests that a much older demographic shift, such as regional differences in the effective sizes of koala populations supported by climatic refugia during the Pleistocene glaciations, may underly this phenomenon (Tsangaras et al., 2012, Neaves et al., 2016, Lott et al., 2022). Interestingly, multilevel mixed-effects linear models failed to detect any relationship between the scale of key threatening processes (e.g., habitat loss) and the level of genetic diversity in NSW koala populations. This highlights the importance of directly assessing genetic diversity when developing risk management frameworks as the genetic health of populations clearly cannot be inferred or predicted from other observable features of the environment.

### 4.4 Management Implications and Directions for Future Research

Despite near universal public support (Brown et al., 2018; Fielding et al., 2022), ongoing attention from the scientific community, and unprecedented financial investment by both the State and Federal governments, koala populations are declining across large parts of NSW. As the rate at which anthropogenic processes destroy or irreversibly alter natural habitats continues to accelerate, the development of strategies for facilitating gene flow between small, fragmented populations will be integral to successful conservation efforts. In the absence of natural dispersal corridors, such strategies are increasingly likely to take the form of translocations. While there are numerous well documented benefits of wildlife translocations, ongoing controversy surrounds their use as a tool for threatened species management (Liddell et al., 2021). Common criticisms include a lack of clear program objectives, poor follow-up monitoring, concerns that translocations do not address, and may even legitimise, the processes that drive local extinctions (e.g., habitat loss), and fears of outbreeding depression caused by the movement of individuals between genetically dissimilar populations (i.e., genetic rescue) (Fischer and Lindenmayer, 2000; Germano et al., 2015; Dresser et al., 2017). Fortunately, there is a growing body of evidence that the risks of outbreeding depression are often exaggerated, and that any negative consequences will typically only persist for a few generations, if they manifest at all (Frankham et al. 2015; Ralls et al. 2020). Furthermore, widespread evidence of past admixture between major genetic clusters of koalas, coupled with the lack of fixed genetic differences between the ARKS, suggests that genetic rescue is a viable conservation strategy for this species. However, there are other concerns about koala translocations that cannot be so easily dismissed. The National Recovery Plan for the Koala 2022 repeatedly acknowledges that translocations are likely to be important for the long-term conservation of this species, while the 2022 NSW Koala Strategy includes the explicit goal of facilitating up to eight translocation projects by 2026. Despite this, there are currently no nationally recognised guidelines for either implementing or critically evaluating the success of koala translocations, and the identification of genetically and ecologically meaningful management divisions remains an ongoing challenge. The results of this study indicate that neither the ARKS nor the derived populations referenced in the NSW Koala Strategy entirely reflect the contemporary distribution of genetic diversity across the State’s koala populations. Most of the major genetic clusters were found to span multiple ARKS, while the Far north-east Hinterland ARKS appeared to contain two genetically distinct groups (Cluster 1 and 2). This is not to suggest that the approach of using koala occurrence records, or data on ecological threats and geographic barriers, to define management divisions lacks merit, simply that care must be taken to ensure that translocation decisions based on these frameworks do not inadvertently restrict gene flow between populations and regions that, historically, are likely to have been interconnected. Similarly, the decision in the NSW Koala Strategy to further divide the Bungonia ARKS into three subregions cannot be justified based on the current genetic data. Koalas across southern NSW represent a single major genetic cluster and display lower levels of diversity on average than their conspecifics further north. Artificially imposing additional divisions on populations that are already relatively small and fragmented may only serve to accelerate the erosion of this diversity. Consequently, the use of translocations to promote gene flow between ARKS that represent the same major genetic clusters should be encouraged where possible, with the understanding that precautions must also be taken to reduce the possibility of negative nongenetic effects caused by the introduction of pathogens (Woodford and Rossiter, 1993; Kock et al., 2010; Dalziel et al., 2017) or the modification of existing host-parasite dynamics (Lott et al., 2012; Aiello et al. 2014; Lott et al., 2015a; Lott et al., 2015b; Lott et al., 2018; Dunlop and Watson, 2022). The movement of individuals between parapatric genetic clusters should also not be ruled out, although the net genetic effects of translocations between specific populations cannot be predicted without further research. The Narrandera ARKS represents one region where the long-term consequences of translocation between genetically divergent koalas could be studied, as this population is a product of admixture between founders from Victoria and north-east NSW/south-east Queensland; two groups of koalas that would not be expected to come into contact naturally (Parsons, 1990; Menkhorst, 2008). Future sample collection efforts targeting regions or management divisions which are currently data deficient will also be critical for determining the precise boundaries between genetically divergent koala populations, particularly those that do not appear to be strongly associated with known biogeographic barriers.

It is important to note that conservation planning generally places a greater emphasis on population-level variation than individual genetic diversity metrics (Avise, 2008; Hoban, 2018; Liddell et al., 2020). However, our data demonstrate that most of the remaining genetic variation to be found in NSW koalas is distributed between individuals rather than among management divisions, or even the major genetic clusters. This strongly suggests that each koala in NSW represents an important reservoir of genetic diversity and evolutionary potential. To enhance conservation outcomes, it is therefore vital that stakeholders reduce koala mortality rates across the entire State, while simultaneously maintaining habitat connectivity and gene flow between as many surviving populations as possible. Given the key knowledge gaps that persist for many koala populations, particularly in southern and western NSW where the species is patchily distributed and detectability is proportionately low, it is highly probable that the development of intensive evidence-based management actions which target specific groups of koalas will ultimately prove impossible. Achieving meaningful conservation outcomes will therefore require the implementation of more robust legislation and management frameworks which address the root causes of ongoing koala population declines (particularly drought and habitat loss). Where existing or emerging data do exist to guide more targeted interventions, the authors recommend that policy makers, land managers, and other stakeholders prioritise the protection of populations which are at the most immediate risk of extinction, rather than those perceived to be more “valuable” based on metrics such as the level or type of genetic diversity that they represent. Our results clearly demonstrate that the loss of any koalas or populations represents a potentially critical reduction of genetic diversity for this iconic Australian marsupial.

## Abbreviations

ACT: Australian Capital Territory
AMOVA: analysis of molecular variance
ARKS: areas of regional koala significance
CLUMPAK: cluster Markov packager across K
DAPC: discriminate analysis of principal components
DArT: Diversity Arrays Technology
EPBC Act: Commonwealth Environment Protection and Biodiversity Conservation Act 1999
GDR: Great Dividing Range
HL: homozygosity by locus
HWE: Hardy–Weinberg equilibrium
IR: internal relatedness
KMA: koala management area
NSW: New South Wales
QLD: Queensland
SA: South Australia
SNP: single nucleotide polymorphism
VIC: Victoria

## Acknowledgements

Funding for this project was provided by the New South Wales Government under the New South Wales Koala Strategy (Grant Agreement No. KR_2019_01), and the Australian Museum Foundation. Samples sourced from living koalas were collected using methods approved under the Australian Museum Animal Care and Ethics Committee (Permit numbers: 11-03, 15_ 05), the New South Wales Director General Animal Ethics Committee (Animal Research Authority approval number: TRIM13/349), the New South Wales Office of Environment and Heritage Scientific License (Permit numbers: SL100280 and SL101687), and the Department of Environment and Primary Industries Animal Ethics Committee (Permit number: 14.14). The authors would like to thank Karen Bettink, Zaiga Deist, James Fitzgerald, Sean Fitzgibbon, Cheyne Flanagan, Sara Goodwin, Bronwyn Houlden, Ros Irwin, Rhonda James, Julie Jennings, Rhonda Pascoe, Ruth Thompson, and Richard Woodman for their assistance with sample collection.

