## Supplemental Information for "Reversing the decline of threatened koala (*Phascolarctos cinereus*) populations in New South Wales: Using genomics to define meaningful conservation goals"

Table of Contents:

| Supplementary material S1 – Sample details  Table S1.1 | Page 2-10 |
| --- | --- |
| Table S1.2 | Page 11-17 |
| Supplementary material S2 – Methods  Figure S2.1 | Page 18 |
| Figure S2.2 | Page 18 |
| Supplementary material S3 - Analysis tables & figures  Table S3.1 | Page 19 |

**Supplementary material S1 – Sample details**

**Table S1.1** Location information for all 314 koala samples genotyped in this study. Where the names of the 50 populations identified in the New South Wales (NSW) Koala Strategy 2022 differ from the corresponding Areas of Regional Koala Significance (ARKS), the former are provided in brackets.

| **Sample ID** | **ARKS** | **Latitude** | **Longitude** | **Major Genetic Cluster**  **(DAPC & STRUCTURE)** |
| --- | --- | --- | --- | --- |
| K_DArTSeq_001 | Armidale | -30.5218 | 151.6494 | Cluster 3 |
| K_DArTSeq_002 | Armidale | -30.4692 | 151.6385 | Cluster 3 |
| K_DArTSeq_003 | Armidale | -30.5128 | 151.6040 | Cluster 3 |
| K_DArTSeq_004 | Armidale | -30.5019 | 151.6463 | Cluster 3 |
| K_DArTSeq_005 | Armidale | -30.4601 | 151.5989 | Cluster 3 |
| K_DArTSeq_006 | Armidale | -30.3130 | 151.6902 | Cluster 3 |
| K_DArTSeq_007 | Armidale | -30.7580 | 151.4516 | Cluster 3 |
| K_DArTSeq_008 | Armidale | -30.4900 | 151.6410 | Cluster 3 |
| K_DArTSeq_009 | Armidale | -30.4637 | 151.1710 | Cluster 3 |
| K_DArTSeq_010 | Armidale | -30.4286 | 151.6593 | Cluster 3 |
| K_DArTSeq_011 | Armidale | -30.6439 | 151.4783 | Cluster 3 |
| K_DArTSeq_012 | Armidale | -30.5410 | 151.7019 | Cluster 3 |
| K_DArTSeq_013 | Armidale | -30.6415 | 151.4916 | Cluster 3 |
| K_DArTSeq_014 | Armidale | -30.6415 | 151.4916 | Cluster 3 |
| K_DArTSeq_015 | Armidale | -30.6415 | 151.4916 | Cluster 3 |
| K_DArTSeq_016 | Armidale | -30.6084 | 151.2251 | Cluster 3 |
| K_DArTSeq_017 | Armidale | -30.4900 | 151.6844 | Cluster 3 |
| K_DArTSeq_018 | Armidale | -30.3361 | 151.6628 | Cluster 3 |
| K_DArTSeq_019 | Armidale | -30.5199 | 151.5193 | Cluster 3 |
| K_DArTSeq_020 | Armidale | -30.4900 | 151.6844 | Cluster 3 |
| K_DArTSeq_021 | Armidale | -30.4900 | 151.6844 | Cluster 3 |
| K_DArTSeq_022 | Armidale | -30.6208 | 151.3417 | Cluster 3 |
| K_DArTSeq_023 | Armidale | -30.4743 | 151.6412 | Cluster 3 |
| K_DArTSeq_024 | Armidale | -30.4798 | 151.6194 | Cluster 3 |
| K_DArTSeq_025 | Armidale | -30.3724 | 151.7135 | Cluster 3 |
| K_DArTSeq_026 | Barrington | -32.4838 | 151.7826 | Cluster 3 |
| K_DArTSeq_027 | Barrington | -32.5688 | 151.7981 | Cluster 3 |
| K_DArTSeq_028 | Belmore River (Crescent Head) | -31.1133 | 152.8342 | Cluster 3 |
| K_DArTSeq_029 | Belmore River (Crescent Head) | -30.9111 | 153.0433 | Cluster 3 |
| K_DArTSeq_030 | Belmore River (Crescent Head) | -31.2419 | 152.9028 | Cluster 3 |
| K_DArTSeq_031 | Belmore River (Crescent Head) | -30.8969 | 153.0438 | Cluster 3 |
| K_DArTSeq_032 | Broadwater | -29.0914 | 153.3978 | Cluster 1 |
| K_DArTSeq_033 | Broadwater | -29.0089 | 153.3975 | Cluster 1 |
| K_DArTSeq_034 | Broadwater | -29.0908 | 153.3697 | Cluster 1 |
| K_DArTSeq_035 | Broadwater | -29.0894 | 153.3722 | Cluster 1 |
| K_DArTSeq_036 | Broadwater | -29.0894 | 153.3722 | Cluster 1 |
| K_DArTSeq_037 | Broadwater | -29.1017 | 153.4300 | Cluster 1 |
| K_DArTSeq_038 | Broadwater | -29.0894 | 153.3722 | Cluster 1 |
| K_DArTSeq_039 | Broadwater | -29.0894 | 153.3722 | Cluster 1 |
| K_DArTSeq_040 | Broadwater | -29.0147 | 153.4336 | Cluster 1 |
| K_DArTSeq_041 | Broadwater | -29.0947 | 153.3964 | Cluster 1 |
| K_DArTSeq_042 | Broadwater | -29.0119 | 153.3958 | Cluster 1 |
| K_DArTSeq_043 | Bungonia (South West Sydney) | -34.2733 | 150.6431 | Cluster 5 |
| K_DArTSeq_044 | Bungonia (South West Sydney) | -34.1361 | 150.8233 | Cluster 5 |
| K_DArTSeq_045 | Bungonia (South West Sydney) | -34.1158 | 151.0617 | Cluster 5 |
| K_DArTSeq_046 | Bungonia (South West Sydney) | -33.9714 | 150.9193 | Cluster 5 |
| K_DArTSeq_047 | Bungonia (South West Sydney) | -34.1865 | 150.7881 | Cluster 5 |
| K_DArTSeq_048 | Bungonia (Southern Highlands) | -34.5321 | 150.5782 | Cluster 5 |
| K_DArTSeq_049 | Bungonia (Southern Highlands) | -34.7937 | 150.1710 | Cluster 5 |
| K_DArTSeq_050 | Bungonia (South-West Sydney) | -34.1228 | 150.8139 | Cluster 5 |
| K_DArTSeq_051 | Bungonia (South-West Sydney) | -34.0736 | 150.8586 | Cluster 5 |
| K_DArTSeq_052 | Bungonia (South-West Sydney) | -34.0736 | 150.8586 | Cluster 5 |
| K_DArTSeq_053 | Bungonia (South-West Sydney) | -34.0736 | 150.8586 | Cluster 5 |
| K_DArTSeq_054 | Bungonia (South-West Sydney) | -34.0736 | 150.8586 | Cluster 5 |
| K_DArTSeq_055 | Bungonia (South-West Sydney) | -34.0736 | 150.8586 | Cluster 5 |
| K_DArTSeq_056 | Bungonia (South-West Sydney) | -34.0736 | 150.8586 | Cluster 5 |
| K_DArTSeq_057 | Bungonia (South-West Sydney) | -34.0736 | 150.8586 | Cluster 5 |
| K_DArTSeq_058 | Bungonia (South-West Sydney) | -34.2369 | 150.6956 | Cluster 5 |
| K_DArTSeq_059 | Coffs Harbour - North Bellingen (Coffs Harbour) | -30.3500 | 153.1000 | Cluster 3 |
| K_DArTSeq_060 | Coffs Harbour - North Bellingen (Coffs Harbour) | -30.4150 | 153.0350 | Cluster 3 |
| K_DArTSeq_061 | Coffs Harbour - North Bellingen (Coffs Harbour) | -30.2500 | 152.8667 | Cluster 3 |
| K_DArTSeq_062 | Coffs Harbour - North Bellingen (Coffs Harbour) | -30.3833 | 153.0333 | Cluster 3 |
| K_DArTSeq_063 | Coffs Harbour - North Bellingen (Coffs Harbour) | -30.3167 | 153.1000 | Cluster 3 |
| K_DArTSeq_064 | Coffs Harbour - North Bellingen (Coffs Harbour) | -30.4069 | 153.0297 | Cluster 3 |
| K_DArTSeq_065 | Coffs Harbour - North Bellingen (Coffs Harbour) | -30.4183 | 153.0242 | Cluster 3 |
| K_DArTSeq_066 | Coffs Harbour - North Bellingen (Coffs Harbour) | -30.4264 | 153.0233 | Cluster 3 |
| K_DArTSeq_067 | Coffs Harbour - North Bellingen (Coffs Harbour) | -30.4125 | 153.0297 | Cluster 3 |
| K_DArTSeq_068 | Coffs Harbour - North Bellingen (Coffs Harbour) | -30.3878 | 153.0394 | Cluster 3 |
| K_DArTSeq_069 | Coffs Harbour - North Bellingen (Coffs Harbour) | -30.4192 | 153.0256 | Cluster 3 |
| K_DArTSeq_070 | Coffs Harbour - North Bellingen (Coffs Harbour) | -30.4328 | 153.0242 | Cluster 3 |
| K_DArTSeq_071 | Coffs Harbour - North Bellingen (Coffs Harbour) | -30.4081 | 153.0267 | Cluster 3 |
| K_DArTSeq_072 | Coffs Harbour - North Bellingen (Coffs Harbour) | -30.4267 | 153.0200 | Cluster 3 |
| K_DArTSeq_073 | Coffs Harbour - North Bellingen (Coffs Harbour) | -30.4150 | 153.0256 | Cluster 3 |
| K_DArTSeq_074 | Coffs Harbour - North Bellingen (Coffs Harbour) | -30.4306 | 153.0625 | Cluster 3 |
| K_DArTSeq_075 | Coffs Harbour - North Bellingen (Coffs Harbour) | -30.4378 | 152.9958 | Cluster 3 |
| K_DArTSeq_076 | Coffs Harbour - North Bellingen (Coffs Harbour) | -30.4042 | 153.0083 | Cluster 3 |
| K_DArTSeq_077 | Coffs Harbour - North Bellingen (Coffs Harbour) | -30.3085 | 153.0853 | Cluster 3 |
| K_DArTSeq_078 | Coffs Harbour - North Bellingen (Coffs Harbour) | -30.0499 | 152.9871 | Cluster 3 |
| K_DArTSeq_079 | Coffs Harbour - North Bellingen (Coffs Harbour) | -30.4453 | 153.0086 | Cluster 3 |
| K_DArTSeq_080 | Coffs Harbour - North Bellingen (Coffs Harbour) | -30.4667 | 153.0500 | Cluster 3 |
| K_DArTSeq_081 | Comboyne | -31.7393 | 152.6908 | Cluster 3 |
| K_DArTSeq_082 | Far north-east | -28.6644 | 153.6050 | Cluster 1 |
| K_DArTSeq_083 | Far north-east | -28.5403 | 153.5433 | Cluster 1 |
| K_DArTSeq_084 | Far north-east | -28.6606 | 153.6144 | Cluster 1 |
| K_DArTSeq_085 | Far north-east | -28.5397 | 153.5436 | Cluster 1 |
| K_DArTSeq_086 | Far north-east | -28.6725 | 153.5528 | Cluster 1 |
| K_DArTSeq_087 | Far north-east | -28.6653 | 153.6105 | Cluster 1 |
| K_DArTSeq_088 | Far north-east Hinterland (Northern Rivers) | -28.9807 | 153.4073 | Cluster 2 |
| K_DArTSeq_089 | Far north-east Hinterland (Northern Rivers) | -28.9786 | 153.4068 | Cluster 1 |
| K_DArTSeq_090 | Far north-east Hinterland (Northern Rivers) | -28.8685 | 153.4418 | Cluster 2 |
| K_DArTSeq_091 | Far north-east Hinterland (Northern Rivers) | -28.9819 | 153.4263 | Cluster 2 |
| K_DArTSeq_092 | Far north-east Hinterland (Northern Rivers) | -28.9825 | 153.4136 | Cluster 2 |
| K_DArTSeq_093 | Far north-east Hinterland (Northern Rivers) | -28.9797 | 153.4066 | Cluster 2 |
| K_DArTSeq_094 | Far north-east Hinterland (Northern Rivers) | -28.9591 | 153.3945 | Cluster 2 |
| K_DArTSeq_095 | Far north-east Hinterland (Northern Rivers) | -28.9850 | 153.4225 | Cluster 2 |
| K_DArTSeq_096 | Far north-east Hinterland (Northern Rivers) | -28.9931 | 153.4368 | Cluster 2 |
| K_DArTSeq_097 | Far north-east Hinterland (Northern Rivers) | -28.9816 | 153.4230 | Cluster 2 |
| K_DArTSeq_098 | Far north-east Hinterland (Northern Rivers) | -28.9931 | 153.4373 | Cluster 2 |
| K_DArTSeq_099 | Far north-east Hinterland (Northern Rivers) | -28.9614 | 153.3942 | Cluster 2 |
| K_DArTSeq_100 | Far north-east Hinterland (Northern Rivers) | -28.9595 | 153.3960 | Cluster 2 |
| K_DArTSeq_101 | Far north-east Hinterland (Northern Rivers) | -28.9825 | 153.4073 | Cluster 2 |
| K_DArTSeq_102 | Far north-east Hinterland (Northern Rivers) | -28.9840 | 153.4304 | Cluster 2 |
| K_DArTSeq_103 | Far north-east Hinterland (Northern Rivers) | -28.9813 | 153.4230 | Cluster 2 |
| K_DArTSeq_104 | Far north-east Hinterland (Northern Rivers) | -28.9403 | 153.4250 | Cluster 2 |
| K_DArTSeq_105 | Far north-east Hinterland (Northern Rivers) | -28.8942 | 153.4469 | Cluster 2 |
| K_DArTSeq_106 | Far north-east Hinterland (Northern Rivers) | -28.8724 | 153.4489 | Cluster 2 |
| K_DArTSeq_107 | Far north-east Hinterland (Northern Rivers) | -28.9285 | 153.4343 | Cluster 2 |
| K_DArTSeq_108 | Far north-east Hinterland (Northern Rivers) | -28.9541 | 153.4570 | Cluster 2 |
| K_DArTSeq_109 | Far north-east Hinterland (Northern Rivers) | -28.9303 | 153.4370 | Cluster 2 |
| K_DArTSeq_110 | Far north-east Hinterland (Northern Rivers) | -28.8640 | 153.4821 | Cluster 2 |
| K_DArTSeq_111 | Far north-east Hinterland (Northern Rivers) | -28.9128 | 153.4378 | Cluster 2 |
| K_DArTSeq_112 | Far north-east Hinterland (Northern Rivers) | -28.9797 | 153.4061 | Cluster 2 |
| K_DArTSeq_113 | Far north-east Hinterland (Northern Rivers) | -28.8689 | 153.4164 | Cluster 2 |
| K_DArTSeq_114 | Far north-east Hinterland (Northern Rivers) | -28.8683 | 153.4166 | Cluster 2 |
| K_DArTSeq_115 | Far north-east Hinterland (Northern Rivers) | -28.8685 | 153.4178 | Cluster 2 |
| K_DArTSeq_116 | Far north-east Hinterland (Northern Rivers) | -28.8685 | 153.4169 | Cluster 2 |
| K_DArTSeq_117 | Far north-east Hinterland (Northern Rivers) | -28.8766 | 153.4451 | Cluster 1 |
| K_DArTSeq_118 | Far north-east Hinterland (Northern Rivers) | -28.8830 | 153.4303 | Cluster 2 |
| K_DArTSeq_119 | Far north-east Hinterland (Northern Rivers) | -28.8000 | 153.3667 | Cluster 2 |
| K_DArTSeq_120 | Far north-east Hinterland (Northern Rivers) | -28.6147 | 152.9572 | Cluster 1 |
| K_DArTSeq_121 | Far north-east Hinterland (Northern Rivers) | -28.8136 | 153.3672 | Cluster 2 |
| K_DArTSeq_122 | Far north-east Hinterland (Northern Rivers) | -28.6975 | 153.3242 | Cluster 1 |
| K_DArTSeq_123 | Far north-east Hinterland (Northern Rivers) | -28.6600 | 153.2850 | Cluster 1 |
| K_DArTSeq_124 | Far north-east Hinterland (Northern Rivers) | -28.6844 | 153.3375 | Cluster 1 |
| K_DArTSeq_125 | Far north-east Hinterland (Northern Rivers) | -28.9492 | 153.3172 | Cluster 2 |
| K_DArTSeq_126 | Far north-east Hinterland (Northern Rivers) | -28.9103 | 153.3064 | Cluster 2 |
| K_DArTSeq_127 | Far north-east Hinterland (Northern Rivers) | -28.7111 | 153.2933 | Cluster 1 |
| K_DArTSeq_128 | Far north-east Hinterland (Northern Rivers) | -28.7319 | 153.4339 | Cluster 1 |
| K_DArTSeq_129 | Far north-east Hinterland (Northern Rivers) | -28.5806 | 153.3745 | Cluster 1 |
| K_DArTSeq_130 | Far north-east Hinterland (Northern Rivers) | -28.6789 | 153.3639 | Cluster 1 |
| K_DArTSeq_131 | Far north-east Hinterland (Northern Rivers) | -28.8219 | 153.3008 | Cluster 2 |
| K_DArTSeq_132 | Far north-east Hinterland (Northern Rivers) | -28.8875 | 153.3003 | Cluster 2 |
| K_DArTSeq_133 | Far north-east Hinterland (Northern Rivers) | -28.8528 | 153.2911 | Cluster 2 |
| K_DArTSeq_134 | Far north-east Hinterland (Northern Rivers) | -28.6833 | 153.3597 | Cluster 1 |
| K_DArTSeq_135 | Far north-east Hinterland (Northern Rivers) | -28.8042 | 153.3411 | Cluster 2 |
| K_DArTSeq_136 | Far north-east Hinterland (Northern Rivers) | -28.3489 | 152.9703 | Cluster 1 |
| K_DArTSeq_137 | Far north-east Hinterland (Northern Rivers) | -28.9108 | 153.3497 | Cluster 2 |
| K_DArTSeq_138 | Far north-east Hinterland (Northern Rivers) | -28.8897 | 153.1908 | Cluster 2 |
| K_DArTSeq_139 | Far north-east Hinterland (Northern Rivers) | -28.8311 | 153.2180 | Cluster 2 |
| K_DArTSeq_140 | Far north-east Hinterland (Northern Rivers) | -28.7456 | 153.2853 | Cluster 1 |
| K_DArTSeq_141 | Far north-east Hinterland (Northern Rivers) | -28.8783 | 153.2450 | Cluster 2 |
| K_DArTSeq_142 | Far north-east Hinterland (Northern Rivers) | -29.0239 | 153.2875 | Cluster 2 |
| K_DArTSeq_143 | Far north-east Hinterland (Northern Rivers) | -28.6600 | 153.1219 | Cluster 1 |
| K_DArTSeq_144 | Far north-east Hinterland (Northern Rivers) | -28.9497 | 153.3186 | Cluster 2 |
| K_DArTSeq_145 | Far north-east Hinterland (Northern Rivers) | -28.4553 | 152.9339 | Cluster 1 |
| K_DArTSeq_146 | Far north-east Hinterland (Northern Rivers) | -28.8900 | 153.3025 | Cluster 2 |
| K_DArTSeq_147 | Far north-east Hinterland (Northern Rivers) | -28.8692 | 153.2397 | Cluster 2 |
| K_DArTSeq_148 | Far north-east Hinterland (Northern Rivers) | -28.8417 | 153.3206 | Cluster 2 |
| K_DArTSeq_149 | Far north-east Hinterland (Northern Rivers) | -28.8594 | 153.3425 | Cluster 2 |
| K_DArTSeq_150 | Far north-east Hinterland (Northern Rivers) | -28.8911 | 153.3206 | Cluster 2 |
| K_DArTSeq_151 | Far north-east Hinterland (Northern Rivers) | -28.8911 | 153.3206 | Cluster 2 |
| K_DArTSeq_152 | Far north-east Hinterland (Northern Rivers) | -28.6758 | 153.4942 | Cluster 2 |
| K_DArTSeq_153 | Far north-east Hinterland (Northern Rivers) | -28.8689 | 153.2828 | Cluster 2 |
| K_DArTSeq_154 | Far north-east Hinterland (Northern Rivers) | -28.6689 | 153.2675 | Cluster 1 |
| K_DArTSeq_155 | Far north-east Hinterland (Northern Rivers) | -28.7389 | 153.3914 | Cluster 1 |
| K_DArTSeq_156 | Far north-east Hinterland (Northern Rivers) | -28.9825 | 153.4220 | Cluster 2 |
| K_DArTSeq_157 | Far north-east Hinterland (Northern Rivers) | -28.6600 | 153.2850 | Cluster 1 |
| K_DArTSeq_158 | Far north-east Hinterland (Northern Rivers) | -28.6444 | 153.4053 | Cluster 1 |
| K_DArTSeq_159 | Far north-east Hinterland (Northern Rivers) | -28.8428 | 153.3031 | Cluster 2 |
| K_DArTSeq_160 | Far north-east Hinterland (Northern Rivers) | -28.7208 | 153.3297 | Cluster 1 |
| K_DArTSeq_161 | Far north-east Hinterland (Northern Rivers) | -28.9108 | 153.4370 | Cluster 2 |
| K_DArTSeq_162 | Far north-east Hinterland (Northern Rivers) | -28.9028 | 153.4833 | Cluster 2 |
| K_DArTSeq_163 | Far north-east Hinterland (Northern Rivers) | -28.9292 | 153.3278 | Cluster 2 |
| K_DArTSeq_164 | Far north-east Hinterland (Northern Rivers) | -28.6603 | 153.4217 | Cluster 1 |
| K_DArTSeq_165 | Far north-east Hinterland (Northern Rivers) | -28.6433 | 152.9992 | Cluster 1 |
| K_DArTSeq_166 | Far north-east Hinterland (Northern Rivers) | -28.9281 | 153.3186 | Cluster 2 |
| K_DArTSeq_167 | Far north-east Hinterland (Northern Rivers) | -28.8136 | 153.3672 | Cluster 2 |
| K_DArTSeq_168 | Far north-east Hinterland (Northern Rivers) | -28.6975 | 153.3242 | Cluster 1 |
| K_DArTSeq_169 | Far north-east Hinterland (Northern Rivers) | -28.6472 | 153.4378 | Cluster 1 |
| K_DArTSeq_170 | Far north-east Hinterland (Northern Rivers) | -28.5806 | 153.3745 | Cluster 1 |
| K_DArTSeq_171 | Far north-east Hinterland (Northern Rivers) | -28.6444 | 153.4053 | Cluster 1 |
| K_DArTSeq_172 | Far north-east Hinterland (Northern Rivers) | -28.8708 | 153.3153 | Cluster 2 |
| K_DArTSeq_173 | Far north-east Hinterland (Northern Rivers) | -28.8311 | 153.2180 | Cluster 2 |
| K_DArTSeq_174 | Far north-east Hinterland (Northern Rivers) | -28.4656 | 152.9183 | Cluster 1 |
| K_DArTSeq_175 | Far north-east Hinterland (Northern Rivers) | -28.8311 | 153.2180 | Cluster 2 |
| K_DArTSeq_176 | Far north-east Hinterland (Northern Rivers) | -28.6606 | 153.4961 | Cluster 2 |
| K_DArTSeq_177 | Far north-east Hinterland (Northern Rivers) | -28.9233 | 153.4339 | Cluster 2 |
| K_DArTSeq_178 | Far north-east Hinterland (Northern Rivers) | -28.6444 | 153.4344 | Cluster 1 |
| K_DArTSeq_179 | Far north-east Hinterland (Northern Rivers) | -28.5692 | 153.0206 | Cluster 1 |
| K_DArTSeq_180 | Far north-east Hinterland (Northern Rivers) | -28.9628 | 153.4547 | Cluster 2 |
| K_DArTSeq_181 | Far north-east Hinterland (Northern Rivers) | -28.3489 | 152.9703 | Cluster 1 |
| K_DArTSeq_182 | Far north-east Hinterland (Northern Rivers) | -28.8725 | 153.0433 | Cluster 1 |
| K_DArTSeq_183 | Far north-east Hinterland (Northern Rivers) | -28.8886 | 153.2997 | Cluster 2 |
| K_DArTSeq_184 | Far north-east Hinterland (Northern Rivers) | -28.8461 | 153.3172 | Cluster 2 |
| K_DArTSeq_185 | Far north-east Hinterland (Northern Rivers) | -28.8461 | 153.3172 | Cluster 2 |
| K_DArTSeq_186 | Far north-east Hinterland (Northern Rivers) | -28.7456 | 153.2853 | Cluster 1 |
| K_DArTSeq_187 | Far north-east Hinterland (Northern Rivers) | -28.6483 | 153.1147 | Cluster 1 |
| K_DArTSeq_188 | Far north-east Hinterland (Northern Rivers) | -28.6167 | 153.0000 | Cluster 1 |
| K_DArTSeq_189 | Gunnedah (Liverpool Plains) | -30.9923 | 150.2369 | Cluster 4 |
| K_DArTSeq_190 | Gunnedah (Liverpool Plains) | -30.9810 | 150.7626 | Cluster 3 |
| K_DArTSeq_191 | Gunnedah (Liverpool Plains) | -31.5753 | 150.4236 | Cluster 4 |
| K_DArTSeq_192 | Gunnedah (Liverpool Plains) | -31.3489 | 150.6469 | Cluster 4 |
| K_DArTSeq_193 | Gunnedah (Liverpool Plains) | -31.1147 | 150.2706 | Cluster 4 |
| K_DArTSeq_194 | Gunnedah (Liverpool Plains) | -30.9887 | 150.2421 | Cluster 3 |
| K_DArTSeq_195 | Gunnedah (Liverpool Plains) | -31.8458 | 149.6832 | Cluster 5 |
| K_DArTSeq_196 | Gunnedah (Liverpool Plains) | -31.3283 | 150.3596 | Cluster 4 |
| K_DArTSeq_197 | Gunnedah (Liverpool Plains) | -31.3280 | 150.3560 | Cluster 4 |
| K_DArTSeq_198 | Gunnedah (Liverpool Plains) | -31.2699 | 150.3612 | Cluster 4 |
| K_DArTSeq_199 | Gunnedah (Liverpool Plains) | -31.1329 | 149.9940 | Cluster 4 |
| K_DArTSeq_200 | Gunnedah (Liverpool Plains) | -31.2362 | 150.1917 | Cluster 4 |
| K_DArTSeq_201 | Gunnedah (Liverpool Plains) | -31.1320 | 149.9971 | Cluster 4 |
| K_DArTSeq_202 | Gunnedah (Liverpool Plains) | -31.1865 | 150.3054 | Cluster 4 |
| K_DArTSeq_203 | Gunnedah (Liverpool Plains) | -31.2756 | 150.2116 | Cluster 4 |
| K_DArTSeq_204 | Gunnedah (Liverpool Plains) | -31.1290 | 150.0011 | Cluster 4 |
| K_DArTSeq_205 | Gunnedah (Liverpool Plains) | -31.2701 | 150.3688 | Cluster 4 |
| K_DArTSeq_206 | Gunnedah (Liverpool Plains) | -31.1886 | 150.3001 | Cluster 4 |
| K_DArTSeq_207 | Gunnedah (Liverpool Plains) | -31.2747 | 150.3803 | Cluster 4 |
| K_DArTSeq_208 | Gunnedah (Liverpool Plains) | -31.2747 | 150.3803 | Cluster 4 |
| K_DArTSeq_209 | Gunnedah (Liverpool Plains) | -31.1350 | 150.0114 | Cluster 4 |
| K_DArTSeq_210 | Gunnedah (Liverpool Plains) | -31.1988 | 150.2756 | Cluster 4 |
| K_DArTSeq_211 | Gunnedah (Liverpool Plains) | -31.1913 | 150.3028 | Cluster 4 |
| K_DArTSeq_212 | Gunnedah (Liverpool Plains) | -31.2013 | 150.2720 | Cluster 4 |
| K_DArTSeq_213 | Gunnedah (Liverpool Plains) | -31.1327 | 150.0137 | Cluster 4 |
| K_DArTSeq_214 | Gunnedah (Liverpool Plains) | -31.1990 | 150.2722 | Cluster 4 |
| K_DArTSeq_215 | Gunnedah (Liverpool Plains) | -31.2617 | 150.3485 | Cluster 4 |
| K_DArTSeq_216 | Gunnedah (Liverpool Plains) | -31.1370 | 150.0058 | Cluster 4 |
| K_DArTSeq_217 | Gunnedah (Liverpool Plains) | -31.1320 | 150.0060 | Cluster 4 |
| K_DArTSeq_218 | Gunnedah (Liverpool Plains) | -31.1729 | 150.3179 | Cluster 4 |
| K_DArTSeq_219 | Gunnedah (Liverpool Plains) | -31.2663 | 150.3765 | Cluster 4 |
| K_DArTSeq_220 | Gunnedah (Liverpool Plains) | -31.2122 | 150.2573 | Cluster 4 |
| K_DArTSeq_221 | Gunnedah (Liverpool Plains) | -31.2262 | 150.1943 | Cluster 4 |
| K_DArTSeq_222 | Gunnedah (Liverpool Plains) | -31.1370 | 150.0058 | Cluster 4 |
| K_DArTSeq_223 | Gunnedah (Liverpool Plains) | -31.1351 | 150.0097 | Cluster 4 |
| K_DArTSeq_224 | Gunnedah (Liverpool Plains) | -31.1336 | 150.0106 | Cluster 4 |
| K_DArTSeq_225 | Gunnedah (Liverpool Plains) | -31.3585 | 150.1221 | Cluster 4 |
| K_DArTSeq_226 | Gunnedah (Liverpool Plains) | -31.1705 | 150.2997 | Cluster 4 |
| K_DArTSeq_227 | Gunnedah (Liverpool Plains) | -31.1349 | 150.0001 | Cluster 4 |
| K_DArTSeq_228 | Gunnedah (Liverpool Plains) | -31.3500 | 150.0831 | Cluster 4 |
| K_DArTSeq_229 | Gunnedah (Liverpool Plains) | -30.9794 | 150.2561 | Cluster 4 |
| K_DArTSeq_230 | Inverell | -29.8939 | 150.6250 | Cluster 3 |
| K_DArTSeq_231 | Inverell | -29.6108 | 150.5450 | Cluster 3 |
| K_DArTSeq_232 | Inverell | -29.6197 | 150.8195 | Cluster 3 |
| K_DArTSeq_233 | Killarney (Narrabri) | -30.3325 | 149.7811 | Cluster 3 |
| K_DArTSeq_234 | Kiwarrak (South Taree) | -31.9968 | 152.4646 | Cluster 3 |
| K_DArTSeq_235 | Narrandera | -34.7333 | 146.5500 | Cluster 5 |
| K_DArTSeq_236 | Narrandera | -34.7333 | 146.5167 | Cluster 5 |
| K_DArTSeq_237 | Narrandera | -34.7333 | 146.5500 | Cluster 5 |
| K_DArTSeq_238 | Narrandera | -34.7333 | 146.5500 | Cluster 1 |
| K_DArTSeq_239 | Narrandera | -34.7500 | 146.5500 | Cluster 5 |
| K_DArTSeq_240 | North Grafton (Grafton) | -29.6819 | 152.9350 | Cluster 3 |
| K_DArTSeq_241 | North Grafton (Grafton) | -29.5703 | 152.7600 | Cluster 3 |
| K_DArTSeq_242 | North Grafton (Grafton) | -29.6083 | 152.8758 | Cluster 3 |
| K_DArTSeq_243 | North Grafton (Grafton) | -29.6072 | 152.8467 | Cluster 3 |
| K_DArTSeq_244 | North Macleay – Nambucca (Nambucca) | -30.7181 | 152.9169 | Cluster 3 |
| K_DArTSeq_245 | Nullica (Eden) | -37.1286 | 149.8219 | Cluster 5 |
| K_DArTSeq_246 | Numeralla  (Southern Tablelands) | -36.1846 | 149.3445 | Cluster 5 |
| K_DArTSeq_247 | Numeralla | -36.1846 | 149.3445 | Cluster 5 |
| K_DArTSeq_248 | (Southern Tablelands) | -36.1846 | 149.3445 | Cluster 5 |
| K_DArTSeq_249 | Numeralla | -36.1846 | 149.3445 | Cluster 5 |
| K_DArTSeq_250 | (Southern Tablelands) | -36.1846 | 149.3445 | Cluster 5 |
| K_DArTSeq_251 | Numeralla | -36.1846 | 149.3445 | Cluster 5 |
| K_DArTSeq_252 | (Southern Tablelands) | -36.1846 | 149.3445 | Cluster 5 |
| K_DArTSeq_253 | Numeralla | -36.1846 | 149.3445 | Cluster 5 |
| K_DArTSeq_254 | (Southern Tablelands) | -36.1846 | 149.3445 | Cluster 5 |
| K_DArTSeq_255 | Numeralla | -36.1846 | 149.3445 | Cluster 5 |
| K_DArTSeq_256 | (Southern Tablelands) | -36.1846 | 149.3445 | Cluster 5 |
| K_DArTSeq_257 | Numeralla | -36.1846 | 149.3445 | Cluster 5 |
| K_DArTSeq_258 | (Southern Tablelands) | -36.1846 | 149.3445 | Cluster 5 |
| K_DArTSeq_259 | Numeralla | -36.1846 | 149.3445 | Cluster 5 |
| K_DArTSeq_260 | (Southern Tablelands) | -36.1846 | 149.3445 | Cluster 5 |
| K_DArTSeq_261 | Numeralla | -36.1846 | 149.3445 | Cluster 5 |
| K_DArTSeq_262 | (Southern Tablelands) | -36.1846 | 149.3445 | Cluster 5 |
| K_DArTSeq_263 | Numeralla | -36.1846 | 149.3445 | Cluster 5 |
| K_DArTSeq_264 | (Southern Tablelands) | -36.1846 | 149.3445 | Cluster 5 |
| K_DArTSeq_265 | Numeralla | -36.1846 | 149.3445 | Cluster 5 |
| K_DArTSeq_266 | (Southern Tablelands) | -36.1846 | 149.3445 | Cluster 5 |
| K_DArTSeq_267 | Numeralla | -36.1846 | 149.3445 | Cluster 5 |
| K_DArTSeq_268 | (Southern Tablelands) | -36.1846 | 149.3445 | Cluster 5 |
| K_DArTSeq_269 | Numeralla | -36.0061 | 149.4011 | Cluster 5 |
| K_DArTSeq_270 | (Southern Tablelands) | -36.1244 | 149.1424 | Cluster 5 |
| K_DArTSeq_271 | Pilliga | -31.2811 | 149.0133 | Cluster 3 |
| K_DArTSeq_272 | Port Macquarie | -31.4678 | 152.9064 | Cluster 3 |
| K_DArTSeq_273 | Port Macquarie | -31.4200 | 152.8672 | Cluster 3 |
| K_DArTSeq_274 | Port Macquarie | -31.4497 | 152.9281 | Cluster 3 |
| K_DArTSeq_275 | Port Macquarie | -31.4544 | 152.9308 | Cluster 3 |
| K_DArTSeq_276 | Port Macquarie | -31.4675 | 152.9147 | Cluster 3 |
| K_DArTSeq_277 | Port Macquarie | -31.4839 | 152.9061 | Cluster 3 |
| K_DArTSeq_278 | Port Macquarie | -31.4322 | 152.8858 | Cluster 3 |
| K_DArTSeq_279 | Port Macquarie | -31.4583 | 152.9128 | Cluster 3 |
| K_DArTSeq_280 | Port Macquarie | -31.4789 | 152.9211 | Cluster 3 |
| K_DArTSeq_281 | Port Macquarie | -31.4522 | 152.8736 | Cluster 3 |
| K_DArTSeq_282 | Port Macquarie | -31.5753 | 152.8231 | Cluster 3 |
| K_DArTSeq_283 | Port Macquarie | -31.4456 | 152.9122 | Cluster 3 |
| K_DArTSeq_284 | Port Macquarie | -31.4394 | 152.8947 | Cluster 3 |
| K_DArTSeq_285 | Port Macquarie | -31.4617 | 152.8747 | Cluster 3 |
| K_DArTSeq_286 | Port Macquarie | -31.4397 | 152.8850 | Cluster 3 |
| K_DArTSeq_287 | Port Macquarie | -31.4678 | 152.9064 | Cluster 3 |
| K_DArTSeq_288 | Port Macquarie | -31.4608 | 152.9236 | Cluster 3 |
| K_DArTSeq_289 | Port Macquarie | -31.4292 | 152.9108 | Cluster 3 |
| K_DArTSeq_290 | Port Macquarie | -31.4642 | 152.8769 | Cluster 3 |
| K_DArTSeq_291 | Port Macquarie | -31.6419 | 152.7939 | Cluster 3 |
| K_DArTSeq_292 | Port Macquarie | -31.4475 | 152.8975 | Cluster 3 |
| K_DArTSeq_293 | Port Stephens | -32.7161 | 152.0698 | Cluster 3 |
| K_DArTSeq_294 | Port Stephens | -32.7323 | 152.1053 | Cluster 3 |
| K_DArTSeq_295 | Port Stephens | -32.7378 | 152.0756 | Cluster 3 |
| K_DArTSeq_296 | Port Stephens | -32.7323 | 152.1049 | Cluster 3 |
| K_DArTSeq_297 | Queen Charlotte’s Creek (Rockley Mount) | -33.5105 | 149.5549 | Cluster 5 |
| K_DArTSeq_298 | Queen Charlotte’s Creek (Rockley Mount) | -33.5167 | 149.2500 | Cluster 5 |
| K_DArTSeq_299 | Queen Charlotte’s Creek (Rockley Mount) | -33.5311 | 149.2551 | Cluster 5 |
| K_DArTSeq_300 | Queen Charlotte’s Creek (Rockley Mount) | -33.5312 | 149.2551 | Cluster 5 |
| K_DArTSeq_301 | Queen Charlotte’s Creek (Rockley Mount) | -33.5312 | 149.2551 | Cluster 5 |
| K_DArTSeq_302 | Queen Charlotte’s Creek (Rockley Mount) | -33.4172 | 149.5773 | Cluster 4 |
| K_DArTSeq_303 | Southern Clarence | -29.6414 | 152.8608 | Cluster 3 |
| K_DArTSeq_304 | Southern Clarence | -29.5942 | 152.9161 | Cluster 3 |
| K_DArTSeq_305 | Tweed Coast | -28.3708 | 153.5600 | Cluster 1 |
| K_DArTSeq_306 | Tweed Coast | -28.4761 | 153.5303 | Cluster 1 |
| K_DArTSeq_307 | Tweed Coast | -28.3083 | 153.5333 | Cluster 1 |
| K_DArTSeq_308 | Tweed Coast | -28.3389 | 153.5333 | Cluster 1 |
| K_DArTSeq_309 | Tweed Ranges | -28.2053 | 153.5219 | Cluster 1 |
| K_DArTSeq_310 | Wang Wauk SF (East Taree) | -32.1231 | 152.3638 | Cluster 3 |
| K_DArTSeq_311 | Wilson River (South Kempsey) | -31.0736 | 152.8828 | Cluster 3 |
| K_DArTSeq_312 | Wollemi NP (Wollemi) | -33.0603 | 150.6975 | Cluster 3 |
| K_DArTSeq_313 | Wollemi NP (Wollemi) | -33.0603 | 150.6975 | Cluster 3 |
| K_DArTSeq_314 | Woodenbong (West Kyogle) | -28.5603 | 152.7980 | Cluster 1 |

**Table S1.2** Untransformed variables used in the multilevel mixed-effects linear model.

| **Sample ID** | **Major Genetic Cluster**  **(DAPC & STRUCTURE)** | **Homozygosity by Locus** | **% Functional Habitat**  **(High & Moderate)** | **% Functional Habitat**  **(Low &**  **Very Low)** | **Human Population Density (per sq. km of land area)** |
| --- | --- | --- | --- | --- | --- |
| K_DArTSeq_001 | Cluster 3 | 0.7679 | 17 | 83 | 3.44 |
| K_DArTSeq_002 | Cluster 3 | 0.7664 | 17 | 83 | 3.44 |
| K_DArTSeq_003 | Cluster 3 | 0.7662 | 17 | 83 | 3.44 |
| K_DArTSeq_004 | Cluster 3 | 0.7629 | 17 | 83 | 3.44 |
| K_DArTSeq_005 | Cluster 3 | 0.7819 | 17 | 83 | 3.44 |
| K_DArTSeq_006 | Cluster 3 | 0.7475 | 17 | 83 | 3.44 |
| K_DArTSeq_007 | Cluster 3 | 0.7885 | 17 | 83 | 3.44 |
| K_DArTSeq_008 | Cluster 3 | 0.7674 | 17 | 83 | 3.44 |
| K_DArTSeq_009 | Cluster 3 | 0.8126 | 17 | 83 | 3.44 |
| K_DArTSeq_010 | Cluster 3 | 0.7885 | 17 | 83 | 3.44 |
| K_DArTSeq_011 | Cluster 3 | 0.7985 | 17 | 83 | 3.44 |
| K_DArTSeq_012 | Cluster 3 | 0.7718 | 17 | 83 | 3.44 |
| K_DArTSeq_013 | Cluster 3 | 0.7494 | 17 | 83 | 3.44 |
| K_DArTSeq_014 | Cluster 3 | 0.7745 | 17 | 83 | 3.44 |
| K_DArTSeq_015 | Cluster 3 | 0.8282 | 17 | 83 | 3.44 |
| K_DArTSeq_016 | Cluster 3 | 0.8019 | 17 | 83 | 3.44 |
| K_DArTSeq_017 | Cluster 3 | 0.7509 | 17 | 83 | 3.44 |
| K_DArTSeq_018 | Cluster 3 | 0.7778 | 17 | 83 | 3.44 |
| K_DArTSeq_019 | Cluster 3 | 0.8144 | 17 | 83 | 3.44 |
| K_DArTSeq_020 | Cluster 3 | 0.7893 | 17 | 83 | 3.44 |
| K_DArTSeq_021 | Cluster 3 | 0.9091 | 17 | 83 | 3.44 |
| K_DArTSeq_022 | Cluster 3 | 0.7983 | 17 | 83 | 3.44 |
| K_DArTSeq_023 | Cluster 3 | 0.7435 | 17 | 83 | 3.44 |
| K_DArTSeq_024 | Cluster 3 | 0.7315 | 17 | 83 | 3.44 |
| K_DArTSeq_025 | Cluster 3 | 0.7616 | 17 | 83 | 3.44 |
| K_DArTSeq_026 | Cluster 3 | 0.9134 | 61 | 39 | 4.30 |
| K_DArTSeq_027 | Cluster 3 | 0.7909 | 61 | 39 | 4.30 |
| K_DArTSeq_028 | Cluster 3 | 0.7695 | 68 | 32 | 8.85 |
| K_DArTSeq_029 | Cluster 3 | 0.9294 | 68 | 32 | 8.85 |
| K_DArTSeq_030 | Cluster 3 | 0.8094 | 68 | 32 | 8.85 |
| K_DArTSeq_031 | Cluster 3 | 0.8480 | 68 | 32 | 8.85 |
| K_DArTSeq_032 | Cluster 1 | 0.7635 | 46 | 54 | 7.70 |
| K_DArTSeq_033 | Cluster 1 | 0.8014 | 46 | 54 | 7.70 |
| K_DArTSeq_034 | Cluster 1 | 0.8544 | 46 | 54 | 7.70 |
| K_DArTSeq_035 | Cluster 1 | 0.8348 | 46 | 54 | 7.70 |
| K_DArTSeq_036 | Cluster 1 | 0.8196 | 46 | 54 | 7.70 |
| K_DArTSeq_037 | Cluster 1 | 0.8065 | 46 | 54 | 7.70 |
| K_DArTSeq_038 | Cluster 1 | 0.7607 | 46 | 54 | 7.70 |
| K_DArTSeq_039 | Cluster 1 | 0.7852 | 46 | 54 | 7.70 |
| K_DArTSeq_040 | Cluster 1 | 0.7682 | 46 | 54 | 7.70 |
| K_DArTSeq_041 | Cluster 1 | 0.7617 | 46 | 54 | 7.70 |
| K_DArTSeq_042 | Cluster 1 | 0.8010 | 46 | 54 | 7.70 |
| K_DArTSeq_043 | Cluster 5 | 0.8409 | 65 | 35 | 21.12 |
| K_DArTSeq_044 | Cluster 5 | 0.6933 | 65 | 35 | 559.10 |
| K_DArTSeq_045 | Cluster 5 | 0.8235 | 65 | 35 | 630.20 |
| K_DArTSeq_046 | Cluster 5 | 0.8316 | 65 | 35 | 10070.00 |
| K_DArTSeq_047 | Cluster 5 | 0.8332 | 65 | 35 | 21.12 |
| K_DArTSeq_048 | Cluster 5 | 0.8125 | 65 | 35 | 19.25 |
| K_DArTSeq_049 | Cluster 5 | 0.8522 | 65 | 35 | 9.79 |
| K_DArTSeq_050 | Cluster 5 | 0.8363 | 65 | 35 | 559.10 |
| K_DArTSeq_051 | Cluster 5 | 0.8815 | 65 | 35 | 559.10 |
| K_DArTSeq_052 | Cluster 5 | 0.8634 | 65 | 35 | 559.10 |
| K_DArTSeq_053 | Cluster 5 | 0.8702 | 65 | 35 | 559.10 |
| K_DArTSeq_054 | Cluster 5 | 0.6316 | 65 | 35 | 559.10 |
| K_DArTSeq_055 | Cluster 5 | 0.8939 | 65 | 35 | 559.10 |
| K_DArTSeq_056 | Cluster 5 | 0.8777 | 65 | 35 | 559.10 |
| K_DArTSeq_057 | Cluster 5 | 0.8268 | 65 | 35 | 559.10 |
| K_DArTSeq_058 | Cluster 5 | 0.8184 | 65 | 35 | 21.12 |
| K_DArTSeq_059 | Cluster 3 | 0.8022 | 64 | 36 | 66.10 |
| K_DArTSeq_060 | Cluster 3 | 0.8323 | 64 | 36 | 66.10 |
| K_DArTSeq_061 | Cluster 3 | 0.7620 | 64 | 36 | 66.10 |
| K_DArTSeq_062 | Cluster 3 | 0.8310 | 64 | 36 | 66.10 |
| K_DArTSeq_063 | Cluster 3 | 0.8333 | 64 | 36 | 66.10 |
| K_DArTSeq_064 | Cluster 3 | 0.7932 | 64 | 36 | 8.20 |
| K_DArTSeq_065 | Cluster 3 | 0.7779 | 64 | 36 | 8.20 |
| K_DArTSeq_066 | Cluster 3 | 0.7620 | 64 | 36 | 8.20 |
| K_DArTSeq_067 | Cluster 3 | 0.7533 | 64 | 36 | 8.20 |
| K_DArTSeq_068 | Cluster 3 | 0.7710 | 64 | 36 | 8.20 |
| K_DArTSeq_069 | Cluster 3 | 0.7747 | 64 | 36 | 8.20 |
| K_DArTSeq_070 | Cluster 3 | 0.7933 | 64 | 36 | 8.20 |
| K_DArTSeq_071 | Cluster 3 | 0.7723 | 64 | 36 | 8.20 |
| K_DArTSeq_072 | Cluster 3 | 0.6990 | 64 | 36 | 8.20 |
| K_DArTSeq_073 | Cluster 3 | 0.7753 | 64 | 36 | 8.20 |
| K_DArTSeq_074 | Cluster 3 | 0.7731 | 64 | 36 | 8.20 |
| K_DArTSeq_075 | Cluster 3 | 0.7775 | 64 | 36 | 8.20 |
| K_DArTSeq_076 | Cluster 3 | 0.7583 | 64 | 36 | 66.10 |
| K_DArTSeq_077 | Cluster 3 | 0.7687 | 64 | 36 | 4.95 |
| K_DArTSeq_078 | Cluster 3 | 0.7805 | 64 | 36 | 8.20 |
| K_DArTSeq_079 | Cluster 3 | 0.7839 | 64 | 36 | 8.20 |
| K_DArTSeq_080 | Cluster 3 | 0.7757 | 64 | 36 | 8.20 |
| K_DArTSeq_081 | Cluster 3 | 0.7861 | 64 | 36 | 9.38 |
| K_DArTSeq_082 | Cluster 1 | 0.7300 | 28 | 72 | 63.13 |
| K_DArTSeq_083 | Cluster 1 | 0.6603 | 28 | 72 | 63.13 |
| K_DArTSeq_084 | Cluster 1 | 0.8048 | 28 | 72 | 63.13 |
| K_DArTSeq_085 | Cluster 1 | 0.7427 | 28 | 72 | 63.13 |
| K_DArTSeq_086 | Cluster 1 | 0.7531 | 28 | 72 | 63.13 |
| K_DArTSeq_087 | Cluster 1 | 0.7490 | 28 | 72 | 63.13 |
| K_DArTSeq_088 | Cluster 2 | 0.7729 | 40 | 60 | 93.25 |
| K_DArTSeq_089 | Cluster 1 | 0.8110 | 40 | 60 | 93.25 |
| K_DArTSeq_090 | Cluster 2 | 0.8090 | 40 | 60 | 93.25 |
| K_DArTSeq_091 | Cluster 2 | 0.7954 | 40 | 60 | 93.25 |
| K_DArTSeq_092 | Cluster 2 | 0.7789 | 40 | 60 | 93.25 |
| K_DArTSeq_093 | Cluster 2 | 0.7355 | 40 | 60 | 93.25 |
| K_DArTSeq_094 | Cluster 2 | 0.8154 | 40 | 60 | 93.25 |
| K_DArTSeq_095 | Cluster 2 | 0.8589 | 40 | 60 | 93.25 |
| K_DArTSeq_096 | Cluster 2 | 0.8015 | 40 | 60 | 93.25 |
| K_DArTSeq_097 | Cluster 2 | 0.7884 | 40 | 60 | 93.25 |
| K_DArTSeq_098 | Cluster 2 | 0.7635 | 40 | 60 | 93.25 |
| K_DArTSeq_099 | Cluster 2 | 0.7750 | 40 | 60 | 93.25 |
| K_DArTSeq_100 | Cluster 2 | 0.7713 | 40 | 60 | 93.25 |
| K_DArTSeq_101 | Cluster 2 | 0.7326 | 40 | 60 | 93.25 |
| K_DArTSeq_102 | Cluster 2 | 0.7718 | 40 | 60 | 93.25 |
| K_DArTSeq_103 | Cluster 2 | 0.7509 | 40 | 60 | 93.25 |
| K_DArTSeq_104 | Cluster 2 | 0.7384 | 40 | 60 | 93.25 |
| K_DArTSeq_105 | Cluster 2 | 0.7503 | 40 | 60 | 93.25 |
| K_DArTSeq_106 | Cluster 2 | 0.7736 | 40 | 60 | 93.25 |
| K_DArTSeq_107 | Cluster 2 | 0.7874 | 40 | 60 | 93.25 |
| K_DArTSeq_108 | Cluster 2 | 0.7560 | 40 | 60 | 93.25 |
| K_DArTSeq_109 | Cluster 2 | 0.7402 | 40 | 60 | 93.25 |
| K_DArTSeq_110 | Cluster 2 | 0.7757 | 40 | 60 | 93.25 |
| K_DArTSeq_111 | Cluster 2 | 0.7476 | 40 | 60 | 93.25 |
| K_DArTSeq_112 | Cluster 2 | 0.7957 | 40 | 60 | 93.25 |
| K_DArTSeq_113 | Cluster 2 | 0.7477 | 40 | 60 | 93.25 |
| K_DArTSeq_114 | Cluster 2 | 0.7931 | 40 | 60 | 93.25 |
| K_DArTSeq_115 | Cluster 2 | 0.7593 | 40 | 60 | 93.25 |
| K_DArTSeq_116 | Cluster 2 | 0.7848 | 40 | 60 | 93.25 |
| K_DArTSeq_117 | Cluster 1 | 0.7257 | 40 | 60 | 93.25 |
| K_DArTSeq_118 | Cluster 2 | 0.7569 | 40 | 60 | 93.25 |
| K_DArTSeq_119 | Cluster 2 | 0.8344 | 40 | 60 | 33.44 |
| K_DArTSeq_120 | Cluster 1 | 0.7462 | 40 | 60 | 2.45 |
| K_DArTSeq_121 | Cluster 2 | 0.7751 | 40 | 60 | 33.44 |
| K_DArTSeq_122 | Cluster 1 | 0.7542 | 40 | 60 | 33.44 |
| K_DArTSeq_123 | Cluster 1 | 0.7369 | 40 | 60 | 33.44 |
| K_DArTSeq_124 | Cluster 1 | 0.7936 | 40 | 60 | 33.44 |
| K_DArTSeq_125 | Cluster 2 | 0.7999 | 40 | 60 | 33.44 |
| K_DArTSeq_126 | Cluster 2 | 0.7633 | 40 | 60 | 33.44 |
| K_DArTSeq_127 | Cluster 1 | 0.7549 | 40 | 60 | 33.44 |
| K_DArTSeq_128 | Cluster 1 | 0.7605 | 40 | 60 | 33.44 |
| K_DArTSeq_129 | Cluster 1 | 0.7374 | 40 | 60 | 33.44 |
| K_DArTSeq_130 | Cluster 1 | 0.7600 | 40 | 60 | 33.44 |
| K_DArTSeq_131 | Cluster 2 | 0.7806 | 40 | 60 | 33.44 |
| K_DArTSeq_132 | Cluster 2 | 0.7640 | 40 | 60 | 33.44 |
| K_DArTSeq_133 | Cluster 2 | 0.7759 | 40 | 60 | 33.44 |
| K_DArTSeq_134 | Cluster 1 | 0.7402 | 40 | 60 | 33.44 |
| K_DArTSeq_135 | Cluster 2 | 0.7789 | 40 | 60 | 33.44 |
| K_DArTSeq_136 | Cluster 1 | 0.7352 | 40 | 60 | 2.45 |
| K_DArTSeq_137 | Cluster 2 | 0.7605 | 40 | 60 | 33.44 |
| K_DArTSeq_138 | Cluster 2 | 0.7954 | 40 | 60 | 33.44 |
| K_DArTSeq_139 | Cluster 2 | 0.7383 | 40 | 60 | 33.44 |
| K_DArTSeq_140 | Cluster 1 | 0.7446 | 40 | 60 | 33.44 |
| K_DArTSeq_141 | Cluster 2 | 0.8087 | 40 | 60 | 33.44 |
| K_DArTSeq_142 | Cluster 2 | 0.7949 | 40 | 60 | 33.44 |
| K_DArTSeq_143 | Cluster 1 | 0.7646 | 40 | 60 | 33.44 |
| K_DArTSeq_144 | Cluster 2 | 0.7713 | 40 | 60 | 33.44 |
| K_DArTSeq_145 | Cluster 1 | 0.7310 | 40 | 60 | 2.45 |
| K_DArTSeq_146 | Cluster 2 | 0.8158 | 40 | 60 | 33.44 |
| K_DArTSeq_147 | Cluster 2 | 0.8350 | 40 | 60 | 33.44 |
| K_DArTSeq_148 | Cluster 2 | 0.8031 | 40 | 60 | 33.44 |
| K_DArTSeq_149 | Cluster 2 | 0.7888 | 40 | 60 | 33.44 |
| K_DArTSeq_150 | Cluster 2 | 0.8670 | 40 | 60 | 33.44 |
| K_DArTSeq_151 | Cluster 2 | 0.8051 | 40 | 60 | 33.44 |
| K_DArTSeq_152 | Cluster 2 | 0.8008 | 40 | 60 | 63.13 |
| K_DArTSeq_153 | Cluster 2 | 0.7422 | 40 | 60 | 33.44 |
| K_DArTSeq_154 | Cluster 1 | 0.7653 | 40 | 60 | 33.44 |
| K_DArTSeq_155 | Cluster 1 | 0.8817 | 40 | 60 | 33.44 |
| K_DArTSeq_156 | Cluster 2 | 0.8069 | 40 | 60 | 93.25 |
| K_DArTSeq_157 | Cluster 1 | 0.7755 | 40 | 60 | 33.44 |
| K_DArTSeq_158 | Cluster 1 | 0.7386 | 40 | 60 | 33.44 |
| K_DArTSeq_159 | Cluster 2 | 0.7921 | 40 | 60 | 33.44 |
| K_DArTSeq_160 | Cluster 1 | 0.7580 | 40 | 60 | 33.44 |
| K_DArTSeq_161 | Cluster 2 | 0.8148 | 40 | 60 | 93.25 |
| K_DArTSeq_162 | Cluster 2 | 0.7940 | 40 | 60 | 93.25 |
| K_DArTSeq_163 | Cluster 2 | 0.7918 | 40 | 60 | 33.44 |
| K_DArTSeq_164 | Cluster 1 | 0.7761 | 40 | 60 | 33.44 |
| K_DArTSeq_165 | Cluster 1 | 0.7384 | 40 | 60 | 2.45 |
| K_DArTSeq_166 | Cluster 2 | 0.7826 | 40 | 60 | 33.44 |
| K_DArTSeq_167 | Cluster 2 | 0.7876 | 40 | 60 | 33.44 |
| K_DArTSeq_168 | Cluster 1 | 0.7842 | 40 | 60 | 33.44 |
| K_DArTSeq_169 | Cluster 1 | 0.7749 | 40 | 60 | 63.13 |
| K_DArTSeq_170 | Cluster 1 | 0.7458 | 40 | 60 | 33.44 |
| K_DArTSeq_171 | Cluster 1 | 0.7762 | 40 | 60 | 33.44 |
| K_DArTSeq_172 | Cluster 2 | 0.8276 | 40 | 60 | 33.44 |
| K_DArTSeq_173 | Cluster 2 | 0.7342 | 40 | 60 | 33.44 |
| K_DArTSeq_174 | Cluster 1 | 0.7827 | 40 | 60 | 2.45 |
| K_DArTSeq_175 | Cluster 2 | 0.7639 | 40 | 60 | 33.44 |
| K_DArTSeq_176 | Cluster 2 | 0.7887 | 40 | 60 | 63.13 |
| K_DArTSeq_177 | Cluster 2 | 0.7438 | 40 | 60 | 93.25 |
| K_DArTSeq_178 | Cluster 1 | 0.7752 | 40 | 60 | 63.13 |
| K_DArTSeq_179 | Cluster 1 | 0.7572 | 40 | 60 | 2.45 |
| K_DArTSeq_180 | Cluster 2 | 0.7683 | 40 | 60 | 93.25 |
| K_DArTSeq_181 | Cluster 1 | 0.7317 | 40 | 60 | 2.45 |
| K_DArTSeq_182 | Cluster 1 | 0.7042 | 40 | 60 | 7.70 |
| K_DArTSeq_183 | Cluster 2 | 0.7663 | 40 | 60 | 33.44 |
| K_DArTSeq_184 | Cluster 2 | 0.7767 | 40 | 60 | 33.44 |
| K_DArTSeq_185 | Cluster 2 | 0.7961 | 40 | 60 | 33.44 |
| K_DArTSeq_186 | Cluster 1 | 0.7432 | 40 | 60 | 33.44 |
| K_DArTSeq_187 | Cluster 1 | 0.7626 | 40 | 60 | 33.44 |
| K_DArTSeq_188 | Cluster 1 | 0.7690 | 40 | 60 | 2.45 |
| K_DArTSeq_189 | Cluster 4 | 0.8512 | 4 | 96 | 2.54 |
| K_DArTSeq_190 | Cluster 3 | 0.7787 | 4 | 96 | 6.32 |
| K_DArTSeq_191 | Cluster 4 | 0.8024 | 4 | 96 | 2.54 |
| K_DArTSeq_192 | Cluster 4 | 0.8108 | 4 | 96 | 1.55 |
| K_DArTSeq_193 | Cluster 4 | 0.8679 | 4 | 96 | 2.54 |
| K_DArTSeq_194 | Cluster 3 | 0.7954 | 4 | 96 | 2.54 |
| K_DArTSeq_195 | Cluster 5 | 0.8109 | 4 | 96 | 0.74 |
| K_DArTSeq_196 | Cluster 4 | 0.8195 | 4 | 96 | 2.54 |
| K_DArTSeq_197 | Cluster 4 | 0.8423 | 4 | 96 | 2.54 |
| K_DArTSeq_198 | Cluster 4 | 0.8443 | 4 | 96 | 2.54 |
| K_DArTSeq_199 | Cluster 4 | 0.8778 | 4 | 96 | 2.54 |
| K_DArTSeq_200 | Cluster 4 | 0.8359 | 4 | 96 | 2.54 |
| K_DArTSeq_201 | Cluster 4 | 0.8631 | 4 | 96 | 2.54 |
| K_DArTSeq_202 | Cluster 4 | 0.8845 | 4 | 96 | 2.54 |
| K_DArTSeq_203 | Cluster 4 | 0.8370 | 4 | 96 | 2.54 |
| K_DArTSeq_204 | Cluster 4 | 0.8769 | 4 | 96 | 2.54 |
| K_DArTSeq_205 | Cluster 4 | 0.8452 | 4 | 96 | 2.54 |
| K_DArTSeq_206 | Cluster 4 | 0.8569 | 4 | 96 | 2.54 |
| K_DArTSeq_207 | Cluster 4 | 0.8840 | 4 | 96 | 2.54 |
| K_DArTSeq_208 | Cluster 4 | 0.8212 | 4 | 96 | 2.54 |
| K_DArTSeq_209 | Cluster 4 | 0.8397 | 4 | 96 | 2.54 |
| K_DArTSeq_210 | Cluster 4 | 0.8509 | 4 | 96 | 2.54 |
| K_DArTSeq_211 | Cluster 4 | 0.8401 | 4 | 96 | 2.54 |
| K_DArTSeq_212 | Cluster 4 | 0.8170 | 4 | 96 | 2.54 |
| K_DArTSeq_213 | Cluster 4 | 0.8646 | 4 | 96 | 2.54 |
| K_DArTSeq_214 | Cluster 4 | 0.8076 | 4 | 96 | 2.54 |
| K_DArTSeq_215 | Cluster 4 | 0.8450 | 4 | 96 | 2.54 |
| K_DArTSeq_216 | Cluster 4 | 0.8618 | 4 | 96 | 2.54 |
| K_DArTSeq_217 | Cluster 4 | 0.8568 | 4 | 96 | 2.54 |
| K_DArTSeq_218 | Cluster 4 | 0.8307 | 4 | 96 | 2.54 |
| K_DArTSeq_219 | Cluster 4 | 0.8439 | 4 | 96 | 2.54 |
| K_DArTSeq_220 | Cluster 4 | 0.8256 | 4 | 96 | 2.54 |
| K_DArTSeq_221 | Cluster 4 | 0.8785 | 4 | 96 | 2.54 |
| K_DArTSeq_222 | Cluster 4 | 0.8320 | 4 | 96 | 2.54 |
| K_DArTSeq_223 | Cluster 4 | 0.8683 | 4 | 96 | 2.54 |
| K_DArTSeq_224 | Cluster 4 | 0.8420 | 4 | 96 | 2.54 |
| K_DArTSeq_225 | Cluster 4 | 0.8507 | 4 | 96 | 2.54 |
| K_DArTSeq_226 | Cluster 4 | 0.8633 | 4 | 96 | 2.54 |
| K_DArTSeq_227 | Cluster 4 | 0.8545 | 4 | 96 | 2.54 |
| K_DArTSeq_228 | Cluster 4 | 0.8261 | 4 | 96 | 2.54 |
| K_DArTSeq_229 | Cluster 4 | 0.8272 | 4 | 96 | 2.54 |
| K_DArTSeq_230 | Cluster 3 | 0.7935 | 5 | 95 | 1.89 |
| K_DArTSeq_231 | Cluster 3 | 0.7856 | 5 | 95 | 1.89 |
| K_DArTSeq_232 | Cluster 3 | 0.7512 | 5 | 95 | 1.89 |
| K_DArTSeq_233 | Cluster 3 | 0.8386 | 5 | 95 | 1.00 |
| K_DArTSeq_234 | Cluster 3 | 0.7859 | 50 | 50 | 9.38 |
| K_DArTSeq_235 | Cluster 5 | 0.8406 | 0 | 100 | 1.42 |
| K_DArTSeq_236 | Cluster 5 | 0.7977 | 0 | 100 | 1.42 |
| K_DArTSeq_237 | Cluster 5 | 0.8390 | 0 | 100 | 1.42 |
| K_DArTSeq_238 | Cluster 1 | 0.7969 | 0 | 100 | 1.42 |
| K_DArTSeq_239 | Cluster 5 | 0.7695 | 0 | 100 | 1.42 |
| K_DArTSeq_240 | Cluster 3 | 0.8172 | 29 | 71 | 14.30 |
| K_DArTSeq_241 | Cluster 3 | 0.7563 | 29 | 71 | 14.30 |
| K_DArTSeq_242 | Cluster 3 | 0.7791 | 29 | 71 | 14.30 |
| K_DArTSeq_243 | Cluster 3 | 0.7727 | 29 | 71 | 14.30 |
| K_DArTSeq_244 | Cluster 3 | 0.7928 | 57 | 43 | 12.89 |
| K_DArTSeq_245 | Cluster 5 | 0.8600 | 85 | 15 | 5.53 |
| K_DArTSeq_246 | Cluster 5 | 0.8052 | 75 | 25 | 27.29 |
| K_DArTSeq_247 | Cluster 5 | 0.8417 | 75 | 25 | 27.29 |
| K_DArTSeq_248 | Cluster 5 | 0.8153 | 75 | 25 | 27.29 |
| K_DArTSeq_249 | Cluster 5 | 0.8104 | 75 | 25 | 27.29 |
| K_DArTSeq_250 | Cluster 5 | 0.8117 | 75 | 25 | 27.29 |
| K_DArTSeq_251 | Cluster 5 | 0.8325 | 75 | 25 | 27.29 |
| K_DArTSeq_252 | Cluster 5 | 0.8060 | 75 | 25 | 27.29 |
| K_DArTSeq_253 | Cluster 5 | 0.8018 | 75 | 25 | 27.29 |
| K_DArTSeq_254 | Cluster 5 | 0.8151 | 75 | 25 | 27.29 |
| K_DArTSeq_255 | Cluster 5 | 0.8062 | 75 | 25 | 27.29 |
| K_DArTSeq_256 | Cluster 5 | 0.7958 | 75 | 25 | 27.29 |
| K_DArTSeq_257 | Cluster 5 | 0.8080 | 75 | 25 | 27.29 |
| K_DArTSeq_258 | Cluster 5 | 0.8192 | 75 | 25 | 27.29 |
| K_DArTSeq_259 | Cluster 5 | 0.8153 | 75 | 25 | 27.29 |
| K_DArTSeq_260 | Cluster 5 | 0.7868 | 75 | 25 | 27.29 |
| K_DArTSeq_261 | Cluster 5 | 0.8455 | 75 | 25 | 27.29 |
| K_DArTSeq_262 | Cluster 5 | 0.8071 | 75 | 25 | 27.29 |
| K_DArTSeq_263 | Cluster 5 | 0.7962 | 75 | 25 | 27.29 |
| K_DArTSeq_264 | Cluster 5 | 0.7992 | 75 | 25 | 27.29 |
| K_DArTSeq_265 | Cluster 5 | 0.7836 | 75 | 25 | 27.29 |
| K_DArTSeq_266 | Cluster 5 | 0.8144 | 75 | 25 | 27.29 |
| K_DArTSeq_267 | Cluster 5 | 0.8044 | 75 | 25 | 27.29 |
| K_DArTSeq_268 | Cluster 5 | 0.8157 | 75 | 25 | 27.29 |
| K_DArTSeq_269 | Cluster 5 | 0.8164 | 75 | 25 | 27.29 |
| K_DArTSeq_270 | Cluster 5 | 0.8160 | 75 | 25 | 27.29 |
| K_DArTSeq_271 | Cluster 3 | 0.8054 | 22 | 78 | 1.00 |
| K_DArTSeq_272 | Cluster 3 | 0.8050 | 51 | 49 | 23.32 |
| K_DArTSeq_273 | Cluster 3 | 0.7999 | 51 | 49 | 23.32 |
| K_DArTSeq_274 | Cluster 3 | 0.7906 | 51 | 49 | 23.32 |
| K_DArTSeq_275 | Cluster 3 | 0.7819 | 51 | 49 | 23.32 |
| K_DArTSeq_276 | Cluster 3 | 0.7710 | 51 | 49 | 23.32 |
| K_DArTSeq_277 | Cluster 3 | 0.8006 | 51 | 49 | 23.32 |
| K_DArTSeq_278 | Cluster 3 | 0.7959 | 51 | 49 | 23.32 |
| K_DArTSeq_279 | Cluster 3 | 0.8047 | 51 | 49 | 23.32 |
| K_DArTSeq_280 | Cluster 3 | 0.8117 | 51 | 49 | 23.32 |
| K_DArTSeq_281 | Cluster 3 | 0.7995 | 51 | 49 | 23.32 |
| K_DArTSeq_282 | Cluster 3 | 0.7823 | 51 | 49 | 23.32 |
| K_DArTSeq_283 | Cluster 3 | 0.6666 | 51 | 49 | 23.32 |
| K_DArTSeq_284 | Cluster 3 | 0.8018 | 51 | 49 | 23.32 |
| K_DArTSeq_285 | Cluster 3 | 0.7881 | 51 | 49 | 23.32 |
| K_DArTSeq_286 | Cluster 3 | 0.7573 | 51 | 49 | 23.32 |
| K_DArTSeq_287 | Cluster 3 | 0.7953 | 51 | 49 | 23.32 |
| K_DArTSeq_288 | Cluster 3 | 0.7653 | 51 | 49 | 23.32 |
| K_DArTSeq_289 | Cluster 3 | 0.7922 | 51 | 49 | 23.32 |
| K_DArTSeq_290 | Cluster 3 | 0.7770 | 51 | 49 | 23.32 |
| K_DArTSeq_291 | Cluster 3 | 0.7338 | 51 | 49 | 23.32 |
| K_DArTSeq_292 | Cluster 3 | 0.8000 | 51 | 49 | 23.32 |
| K_DArTSeq_293 | Cluster 3 | 0.9025 | 64 | 36 | 21.29 |
| K_DArTSeq_294 | Cluster 3 | 0.9039 | 64 | 36 | 21.29 |
| K_DArTSeq_295 | Cluster 3 | 0.9184 | 64 | 36 | 21.29 |
| K_DArTSeq_296 | Cluster 3 | 0.9167 | 64 | 36 | 21.29 |
| K_DArTSeq_297 | Cluster 5 | 0.8407 | 28 | 68 | 11.52 |
| K_DArTSeq_298 | Cluster 5 | 0.7955 | 28 | 68 | 4.84 |
| K_DArTSeq_299 | Cluster 5 | 0.8143 | 28 | 68 | 4.84 |
| K_DArTSeq_300 | Cluster 5 | 0.8185 | 28 | 68 | 4.84 |
| K_DArTSeq_301 | Cluster 5 | 0.8358 | 28 | 68 | 4.84 |
| K_DArTSeq_302 | Cluster 4 | 0.8135 | 28 | 68 | 11.52 |
| K_DArTSeq_303 | Cluster 3 | 0.7844 | 25 | 75 | 4.95 |
| K_DArTSeq_304 | Cluster 3 | 0.7560 | 25 | 75 | 4.95 |
| K_DArTSeq_305 | Cluster 1 | 0.7585 | 32 | 68 | 75.15 |
| K_DArTSeq_306 | Cluster 1 | 0.7709 | 32 | 68 | 63.13 |
| K_DArTSeq_307 | Cluster 1 | 0.7584 | 32 | 68 | 75.15 |
| K_DArTSeq_308 | Cluster 1 | 0.7723 | 32 | 68 | 75.15 |
| K_DArTSeq_309 | Cluster 1 | 0.7552 | 36 | 64 | 75.15 |
| K_DArTSeq_310 | Cluster 3 | 0.7879 | 67 | 33 | 12.48 |
| K_DArTSeq_311 | Cluster 3 | 0.8013 | 65 | 35 | 8.85 |
| K_DArTSeq_312 | Cluster 3 | 0.8003 | 79 | 21 | 21.29 |
| K_DArTSeq_313 | Cluster 3 | 0.8058 | 79 | 21 | 21.29 |
| K_DArTSeq_314 | Cluster 1 | 0.7384 | 61 | 39 | 2.45 |

**Supplementary material S2 – Methods**

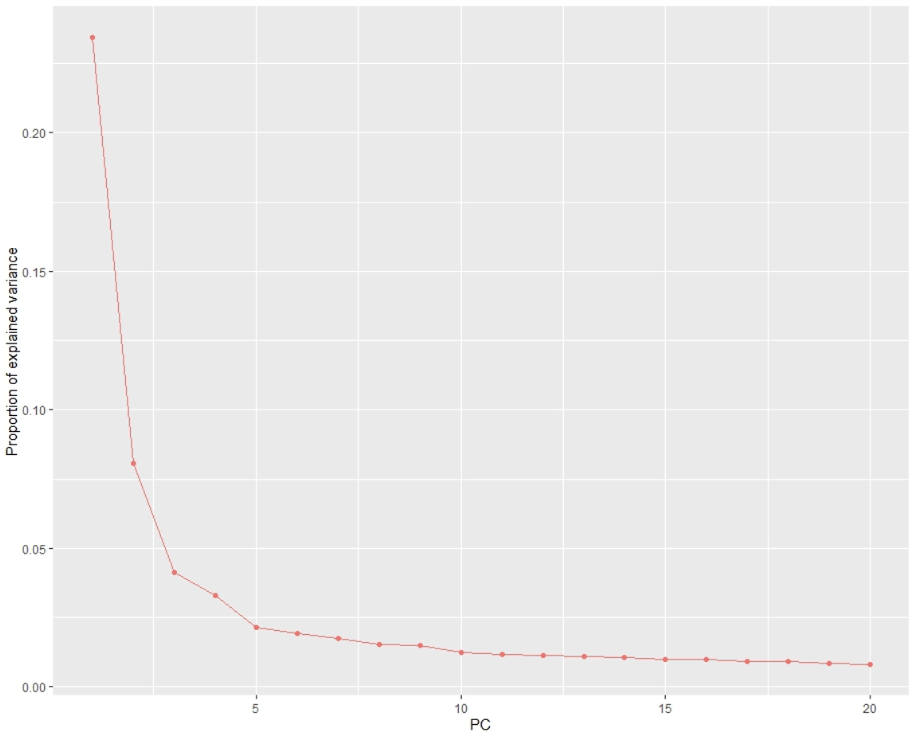

**Figure S2.1** PCAdapt ‘scree plot’ depicting the percentage of variance explained by each PC in decreasing order.

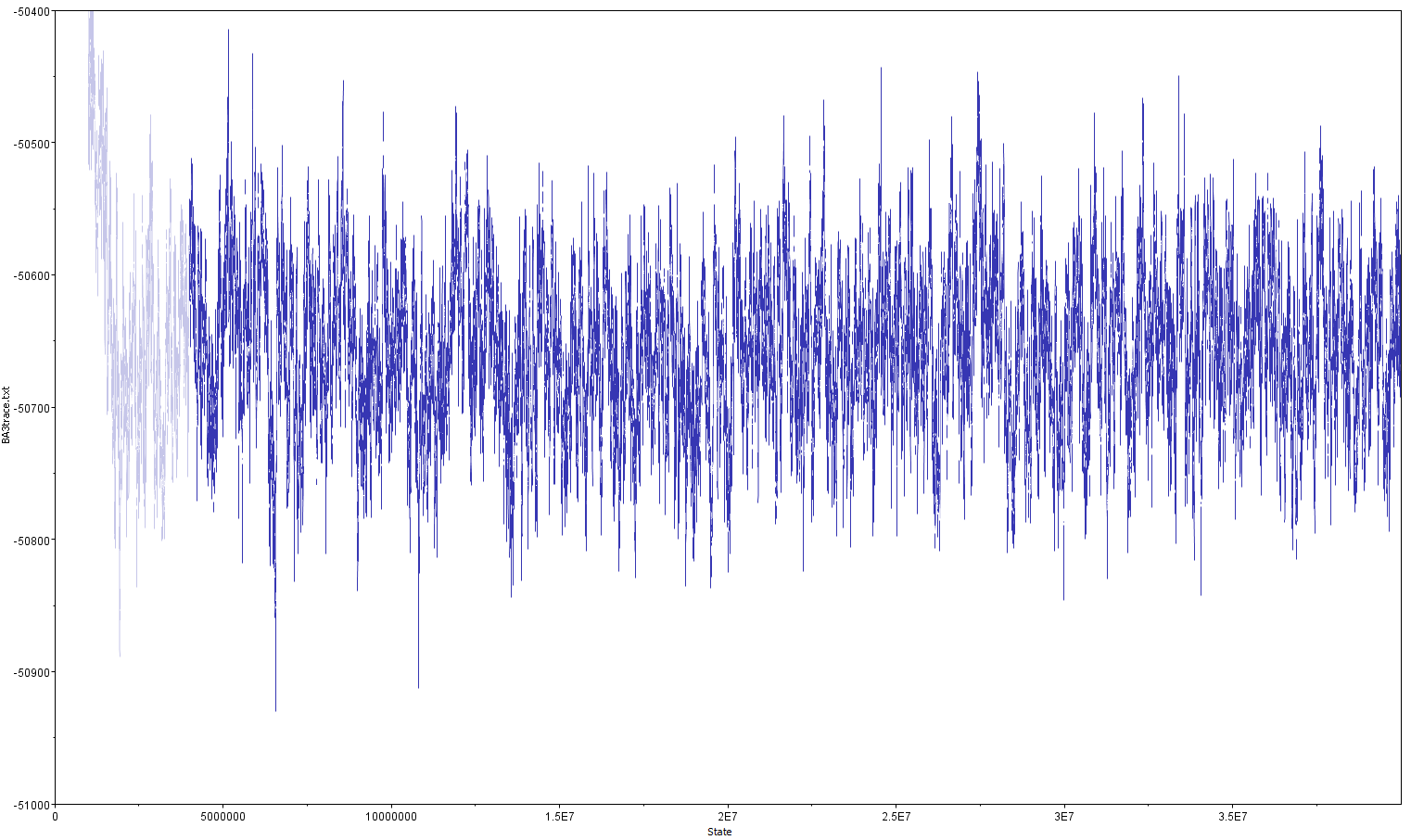

**Figure S2.2** Bayesian posterior parameter trace plot (BayesAss). The x-axis displays the iterations of the MCMC procedures and the y-axis shows the corresponding parameter values. The burn-in iterations are indicated in light blue.

**Supplementary material S3 - Analysis tables & figures**

**Table S3.1** Means and 95% confidence intervals of the posterior distributions for contemporary koala migration rates. Pairwise values significantly different from zero are denoted with an asterisk.

|  | Recipient Population | | | | | | |
| --- | --- | --- | --- | --- | --- | --- | --- |
| Source Population |  | Cluster 1 | Cluster 2 | Cluster 3_  East GDR | Cluster 3_  West GDR | Cluster 4 | Cluster 5 |
|  | Cluster 1 | **0.901* (0.839-0.964)** | 0.250* (0.188-0.312) | 0.011 (-0.010-0.033) | 0.042 (-0.010-0.094) | 0.022 (-0.019-0.063) | 0.024 (-0.003-0.051) |
|  | Cluster 2 | 0.013 (-0.011-0.038) | **0.683* (0.652-0.714)** | 0.011 (-0.010-0.032) | 0.020 (-0.017-0.059) | 0.022 (-0.019-0.063) | 0.008 (-0.007-0.024) |
|  | Cluster 3_  East GDR | 0.058* (0.002-0.113) | 0.017 (-0.014-0.048) | **0.933* (0.886-0.980)** | 0.208* (0.131-0.285) | 0.022 (-0.019-0.063) | 0.009 (-0.008-0.026) |
|  | Cluster 3_  West GDR | 0.013 (-0.012-0.039) | 0.017 (-0.014-0.047) | 0.011 (-0.010-0.032) | **0.687* (0.649-0.725)** | 0.022 (-0.019-0.063) | 0.008 (-0.008-0.024) |
|  | Cluster 4 | 0.013 (-0.012-0.039) | 0.017 (-0.014-0.048) | 0.011 (-0.010-0.011) | 0.021 (-0.018-0.060) | **0.911* (0.655-0.711)** | 0.008 (-0.008-0.024) |
|  | Cluster 5 | 0.013 (-0.012-0.038) | 0.017 (-0.015-0.048) | 0.022 (-0.007-0.052) | 0.021 (-0.018-0.060) | 0.022 (-0.018-0.063) | **0.942* (0.903-0.981)** |
